# Computational Counterfactuals Reveal Non-Additive Audiovisual Semantics in Natural Movie Responses

**DOI:** 10.64898/2026.07.12.738026

**Authors:** Muwei Li

**Affiliations:** Vanderbilt University Institute of Imaging Science, Vanderbilt University Medical Center, Nashville, TN, USA; Department of Radiology and Radiological Sciences, Vanderbilt University Medical Center, Nashville, TN, USA

**Author notes:** Corresponding author. Vanderbilt University Institute of Imaging Science, 1161 21st Ave. S, Medical Center North, AA-1105, Nashville, TN 37232-2310, USA.

**Keywords:** Naturalistic fMRI, Movie Watching, Audiovisual Integration, Semantic Encoding, LLM

## Abstract

Natural audiovisual perception may not be fully captured by decomposing movies into auditory and visual streams. I introduce a computational-counterfactual framework that keeps movie viewing intact while varying only AI-derived descriptions of the same clips. Using 7 Tesla movie fMRI imaging data from 176 participants, I tested whether cortical responses were better predicted by native audiovisual semantics than by a dimension-matched additive reconstruction from audio-only and video-only descriptions. The native model outperformed the matched additive baseline under content-aware purged cross-validation, with strongest gains in auditory, visual, and dorsal attention systems. Representational-similarity, feature-replacement, and content-gating analyses showed that the advantage reflected feature- and network-specific routing linked to coherent audiovisual semantic emergence rather than raw auditory-visual discrepancy. The effect survived stronger temporal purging and repeat-content exclusion, suggesting that intact movie viewing evokes cortical structure aligned with native audiovisual meaning beyond additive unimodal semantics.

## INTRODUCTION

In natural environments, human beings continuously integrate auditory and visual information to construct coherent, time-evolving semantic interpretations of the world^1^. Because real-world understanding depends on the ongoing coordination of multimodal information across time, naturalistic movie-watching paradigms provide a powerful framework for studying this process under conditions of high ecological validity^2,3^. Compared with resting-state or highly simplified task-based designs, naturalistic audiovisual stimuli preserve the continuous, multimodal, and narrative structure of real life while evoking reliable and behaviorally informative cortical responses^4–7^. In this context, a natural extension of classical multisensory logic would be to decompose continuous audiovisual experience by manipulating or removing one sensory modality at a time.

Many early neuroimaging studies operationalized audiovisual integration by comparing cortical responses to combined audiovisual stimulation (AV) with responses to the corresponding auditory-only (A) and visual-only conditions (V), often applying an additive criterion such as AV>A+V to identify superadditive responses^8,9^. However, the validity of this contrast depends on the conditions differing primarily in their sensory inputs, an assumption that is difficult to sustain for continuous naturalistic stimuli, whose sensory, semantic, attentional, and narrative properties covary over time^10–12^. Muting an ongoing movie is therefore not a sensory subtraction alone. Original soundtracks alter gaze location, fixation timing, and the degree to which different viewers attend to the same elements of a scene^13–16^. These attentional differences are consequential for neural measurements because attentional engagement strongly modulates the reliability of cortical responses to naturalistic narratives. Removing sound can also increase cognitive effort, modestly reduce comprehension, and substantially reduce immersion during video viewing^17,18^. Moreover, audio affects the perceptual continuity with which viewers process edits between successive shots^19,20^. Under such conditions, physical deprivation may distort the very representational structure one aims to study. A key unresolved problem, therefore, is how to isolate cortical sensitivity to joint audiovisual meaning without disrupting the natural viewing experience itself.

To address this problem, I reverse the conventional experimental logic. Rather than physically altering the stimulus presented to participants, I keep the complete audiovisual movie fixed, continuous, and naturalistic throughout scanning and vary only the computational description of the stimulus. I term this strategy computational counterfactuals. This strategy holds the sensory and cognitive state of participants constant across model comparisons while asking how the same intact event can be represented through joint audiovisual and separate unimodal accounts. It does not remove the intrinsic covariance among naturalistic stimulus features. Instead, it avoids introducing additional participant-level confounds through physical modality deprivation. Recent advances in computational neuroscience and artificial intelligence make this strategy feasible. Large language models and multimodal artificial intelligence systems can extract rich, high-level semantic features from complex naturalistic audiovisual streams and organize them into psychologically interpretable dimensions^21–23^. My recent work further showed that a multimodal large language model can serve as an automated semantic annotator that links continuous movie content, distributed cortical responses, and cognitive variation through psychologically interpretable dimensions^24^. Together, these developments make it possible to construct alternative computational representations of the same intact naturalistic event without presenting altered stimuli to participants.

I implement this framework by comparing two feature spaces defined over the same movie segments and the same predefined semantic dimensions. The Native Representation comprises joint semantic features extracted directly from intact audiovisual clips. The Matched Additive Representation is a dimension-matched baseline constructed by combining separate semantic estimates generated from the corresponding audio-only and silent video inputs. Crucially, these unimodal inputs are presented only to the artificial intelligence model and never to the participants. The central question is whether cortical activity is better predicted by the native joint representation than by an additive composition of unimodal semantic information. This comparison does not test classical superadditivity of the blood oxygen level-dependent signal. Instead, it tests representational nonadditivity. Specifically, it asks whether intact audiovisual semantics capture predictive cortical structure beyond the structure explained by a matched additive account of unimodal meaning.

To test this hypothesis, I analyze high-quality 7 Tesla movie fMRI data from 176 participants in the Human Connectome Project. I developed a strictly controlled, content-aware purged leave-one-clip-out encoding framework in which all training clips sharing overlapping or homologous narrative content with the test clip are removed, including repeated material presented across different runs. I then compare Native and Matched Additive semantic models that are explicitly aligned across twelve predefined dimensions, thereby ruling out the trivial explanation that any predictive advantage is driven simply by differences in feature dimensionality or gross model capacity.

I hypothesize that cortical activity during natural audiovisual experience contains region-specific and feature-specific predictive information that is more faithfully captured by native joint semantic representations than by matched additive unimodal semantics. Under this view, any Native advantage would not imply a classical claim of blood-oxygen-level-dependent superadditivity but would instead indicate that cortical responses preserve representational structure specific to intact audiovisual meaning. By testing this possibility, my study aims to establish a principled framework for examining multisensory semantic processing under naturalistic conditions without altering the experience of the participants, and to show how multimodal artificial intelligence can be used to probe the organization of complex brain activity at the level of computational representations.

## RESULTS

### Computational counterfactuals compare native and matched additive audiovisual semantics

I used model names that encode both information source and dimensionality: Audio9 and Video8 denote unimodal audio-conditioned and silent-video feature spaces; NativeAV12 denotes 12 intact audiovisual semantic scores; MatchedAdditive12 denotes the dimension-matched counterfactual assembled from unimodal scores; and LateConcat17, ConcatPCA12, and LegacyConcat24 are concatenation and capacity controls. Full feature construction is described in Methods.

To test whether intact audiovisual semantics predicted cortical activity beyond a dimension-matched account assembled from unimodal descriptions, I segmented the continuous movie runs into partially overlapping clips, scored the same movie clips with Gemini under intact audiovisual, audio-only, and silent-video inputs, constructed a 12-dimensional Native audiovisual representation and a 12-dimensional Matched Additive representation, and compared their ridge-encoding performance across 360 cortical parcels in 176 participants. Native multimodal advantage (NMA) was defined as cross-validated R²_NativeAV12_ minus R²_MatchedAdditive12_. Here, R² quantified how well a feature space predicted held-out clip-averaged cortical responses across movie clips for each parcel. Higher R² indicates that clip-to-clip variation in the AI-derived semantic scores explained more clip-to-clip variation in cortical activity. Clip-level prediction used content-aware purged leave-one-clip-out cross-validation with source- excluded nested regularization, and participant inference adjusted for motion, age, and sex. Full procedural and statistical details are provided in the Methods.

The computational counterfactual design held the viewed stimulus and measured brain responses fixed while changing only the representation used to predict those responses (Fig. 1a). The four movie runs initially yielded 293 overlapping clips in 18 presentation blocks. Quality control retained 270 clips in 17 blocks, spanning 13 non-repeated source films and four presentations of one repeated Vimeo source. 28 retained clips belonged to the repeated content (Fig. 1b). The run timeline, block-level clip counts, sample attrition, and day/session/phase-encoding structure are documented in Supplementary Fig. S1a-d.

**Figure 1.**
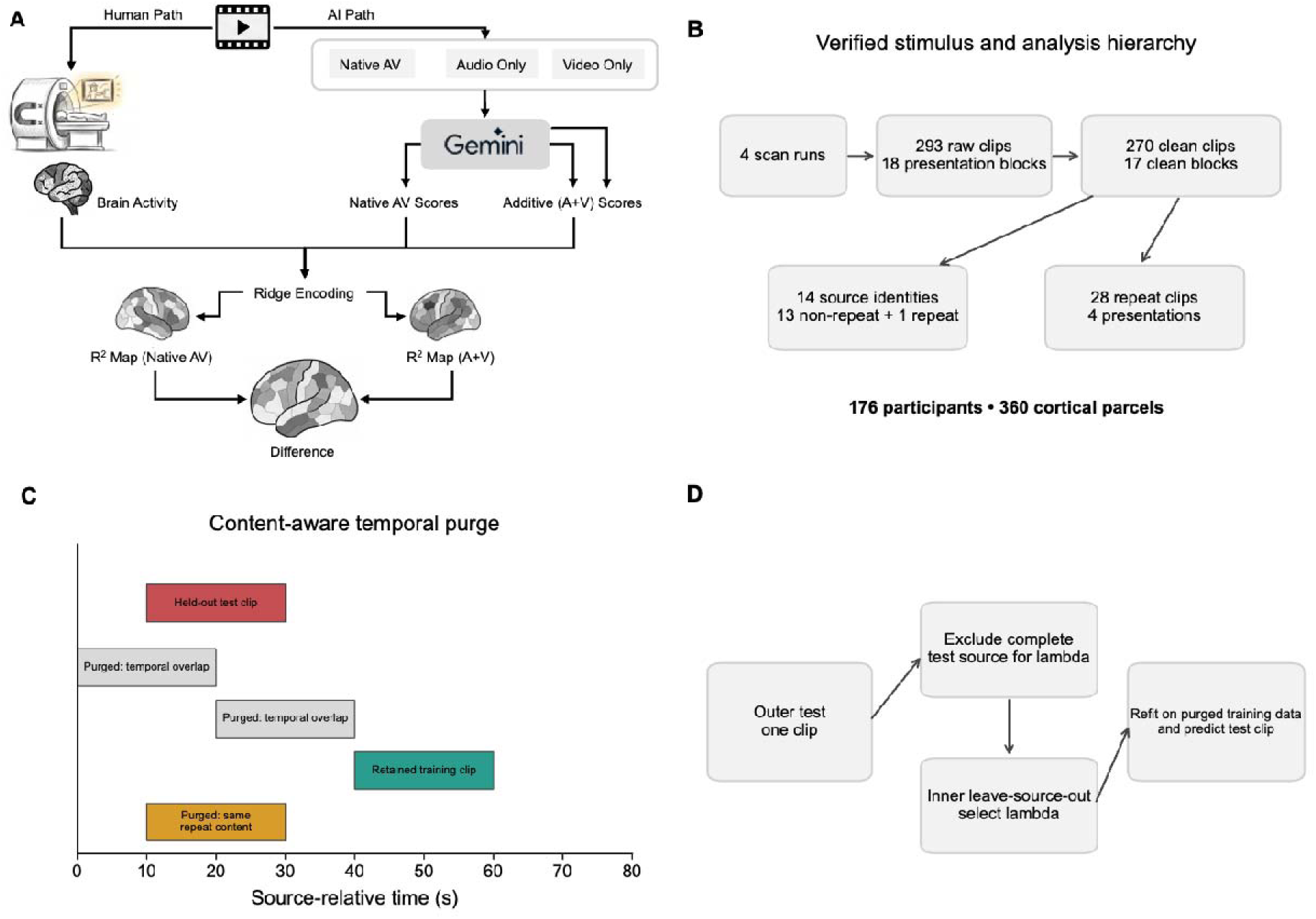
Computational counterfactual design, stimulus hierarchy, and nested cross-validation. (a) The same intact audiovisual clips and observed fMRI responses were used for every model comparison; only the computational stimulus representation changed. NativeAV12 was scored directly from intact audiovisual clips, whereas MatchedAdditive12 was assembled on the same 12 semantic dimensions from audio-only and silent-video scores. (b) Four scan runs yielded 293 raw clips in 18 presentation blocks; quality control retained 270 clips in 17 blocks, 14 source identities (13 non-repeat sources and one repeated source), 28 repeat clips across four presentations, 176 participants, and 360 MMP parcels. (c) In content-aware purged leave-one-clip-out cross-validation, same-source clips with overlapping time intervals were removed from training; the aligned cross-run counterpart of repeated Vimeo content was also removed. Unrelated clips were retained. (d) Ridge lambda was selected by inner leave-source-out validation after excluding the complete outer-test source; the selected model was then refitted on the purged training data and used to predict the held-out clip. Full details are provided in Methods.

NativeAV12 comprised semantic scores extracted directly from each intact audiovisual clip. MatchedAdditive12 used the corresponding silent-video scores for visual-specific dimensions, audio-only scores for auditory-specific dimensions, and the mean of independently obtained audio-only and silent-video scores for five shared abstract dimensions. Thus, the two primary models occupied the same 12 named semantic axes and differed in how those axes were derived, rather than in dimensionality. Dimension-wise Native-Matched correspondence, audio-video correspondence, the correlation structure of Native-minus-Matched residuals, and feature medians confirmed that the spaces were related but not identical (Supplementary Fig. S2a-d).

Because adjacent clips overlapped in time, each outer fold removed same-source training clips whose time intervals overlapped the held-out clip. For the repeated Vimeo content, the aligned cross-run counterpart was also removed, whereas clips from unrelated content remained eligible for training (Fig. 1c). Ridge regularization was selected inside each outer fold by leave- source-out validation that excluded the complete test source, after which the model was refitted on the purged training set and used to predict the held-out clip (Fig. 1d). The distribution of purge counts, model-specific lambda ranges, source-film sample sizes, and the source- excluded selection scheme are shown in Supplementary Fig. S3a-d.

### Native audiovisual semantics improve cortical encoding

Across the seven prespecified feature spaces, NativeAV12 produced the highest cortex-wide group mean cross-validated R² (0.09610), compared with 0.07466 for MatchedAdditive12, a difference of 0.02144 (Fig. 2a). NativeAV12 also exceeded the higher-dimensional LateConcat17 (0.08283), the dimension-matched ConcatPCA12 (0.08183), and LegacyConcat24 (0.07651), as well as the Audio9 (0.02676) and Video8 (0.06858) unimodal models. This ordering argues against a simple explanation based on feature count or generic concatenation. The same model-capacity comparisons are summarized as a dedicated sensitivity analysis in Supplementary Fig. S18a.

**Figure 2.**
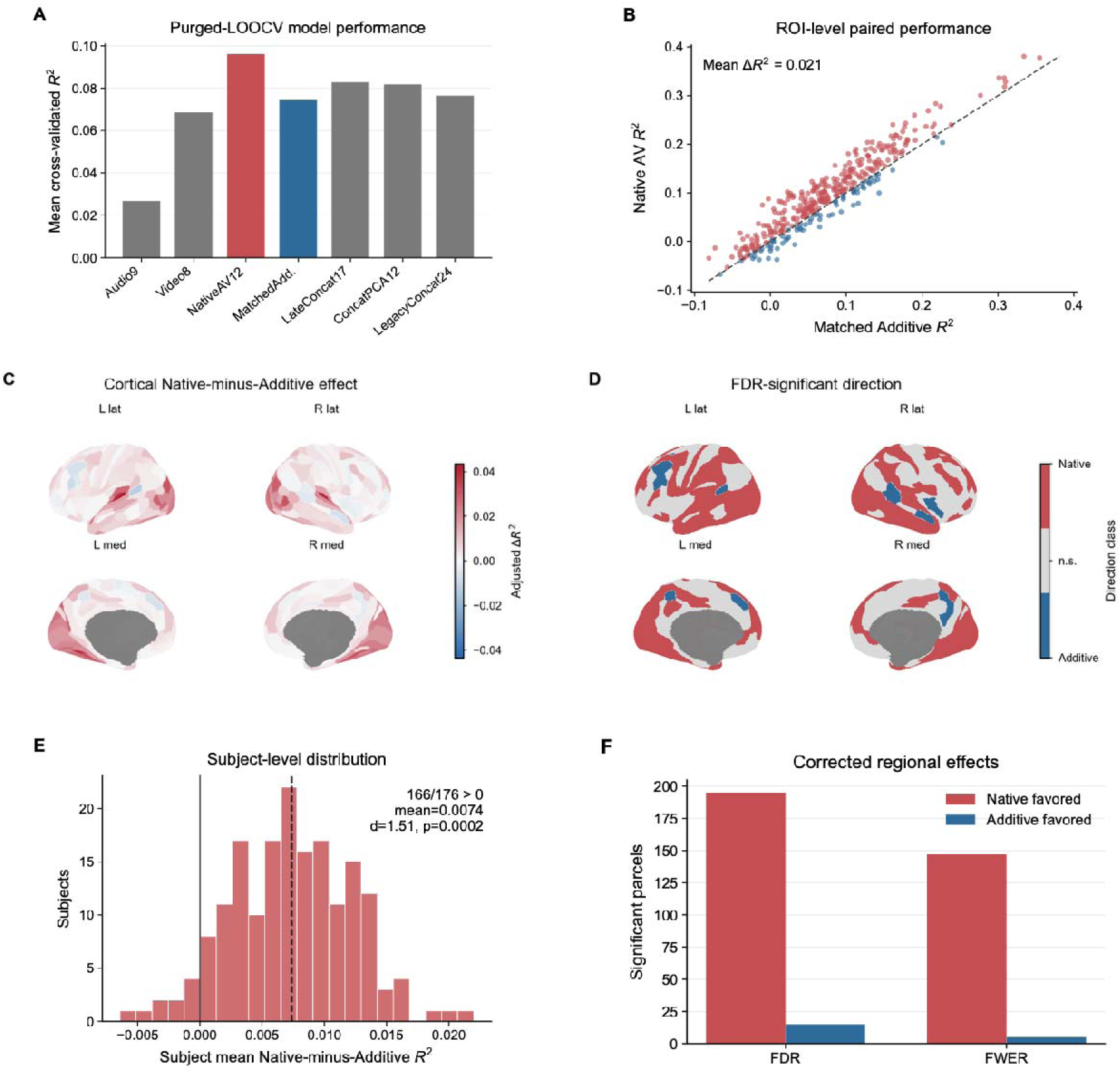
Native audiovisual scoring improves purged cortical encoding. (a) Cortex-wide mean cross-validated R² for seven prespecified feature spaces. Audio9 and Video8 are unimodal models; NativeAV12 and MatchedAdditive12 are the primary dimension-matched comparison; LateConcat17, ConcatPCA12, and LegacyConcat24 control for concatenation strategy and capacity. (b) Paired parcel- level Native and Matched Additive R²; the dashed diagonal denotes equality. (c) Covariate-adjusted Native-minus-Additive R² (NMA) across 360 MMP parcels. (d) Direction of parcels surviving two-sided FDR across 360 tests; red favors Native and blue favors Matched Additive. (e) Distribution of cortex- averaged NMA across 176 participants; the solid vertical line denotes zero and the dashed line the sample mean. The displayed Cohen’s d and two-sided bootstrap p value use participants as the inferential unit after adjustment for motion, age, and sex. (f) Counts of Native- and Additive-favored parcels surviving FDR and max-statistic FWER.

The Native advantage was visible across the paired parcel-wise performance distribution (Fig. 2b), although it was not uniform over cortex. Absolute out-of-sample R² maps for every model, shown on a common scale, localized the different prediction regimes and confirmed that the primary contrast was not driven by a single anomalous baseline map (Supplementary Fig. S4a- g). The participant-adjusted NMA map showed broad positive effects in auditory and visual territories, accompanied by smaller negative effects in a restricted set of association parcels (Fig. 2c). Directional thresholding retained both classes rather than treating the Native advantage as a one-sided whole-cortex effect (Fig. 2d).

At the participant level, after adjustment for motion, age, and sex, the cortex-averaged NMA was 0.007372 (t=19.99, Cohen’s d=1.51, two-sided bootstrap p=0.0002), and 94.3% of participants had a positive cortex-averaged difference (Fig. 2e). Of 360 parcels, 210 survived two-sided false-discovery-rate (FDR) correction and 152 survived max-statistic family-wise-error-rate (FWER) correction. Under FDR, 195 parcels favored Native and 15 favored Matched Additive; under FWER, the corresponding counts were 147 and 5 (Fig. 2f). Complete maps of adjusted effect size, t statistic, Cohen’s d, FWER significance, participant-wise sign consistency, and signed FDR evidence are provided in Supplementary Fig. S5a-f.

The strongest Native-favored parcels included bilateral auditory belt and primary auditory regions, whereas the largest Additive-favored effects included left STV, IFSp, 8C, and 8BM (Supplementary Fig. S6a, b). Network parcel counts and hemispheric counts of significant positive and negative effects showed that this pattern was not reducible to unequal network size or a single hemisphere (Supplementary Fig. S6c, d). Together, these results support a broad but spatially graded Native advantage with reproducible local exceptions.

### Native gain follows a graded network hierarchy and extends to representational geometry

Aggregation within the 12 Cole-Anticevic Brain Network Partition networks revealed significant positive amplitude NMA in 11 networks after two-sided FDR correction; only the orbito-affective network did not survive correction (Fig. 3a). The largest effects occurred in auditory cortex (AUD; NMA=0.03145, d=1.87), secondary visual cortex (VIS2; 0.01849, d=1.76), primary visual cortex (VIS1; 0.01699, d=1.21), and the dorsal-attention network (DAN; 0.01236, d=1.34). Mapping each network estimate back to its constituent parcels emphasized the strong sensory and attention components of the effect rather than confining Native gain to high-level association cortex (Fig. 3b).

**Figure 3.**
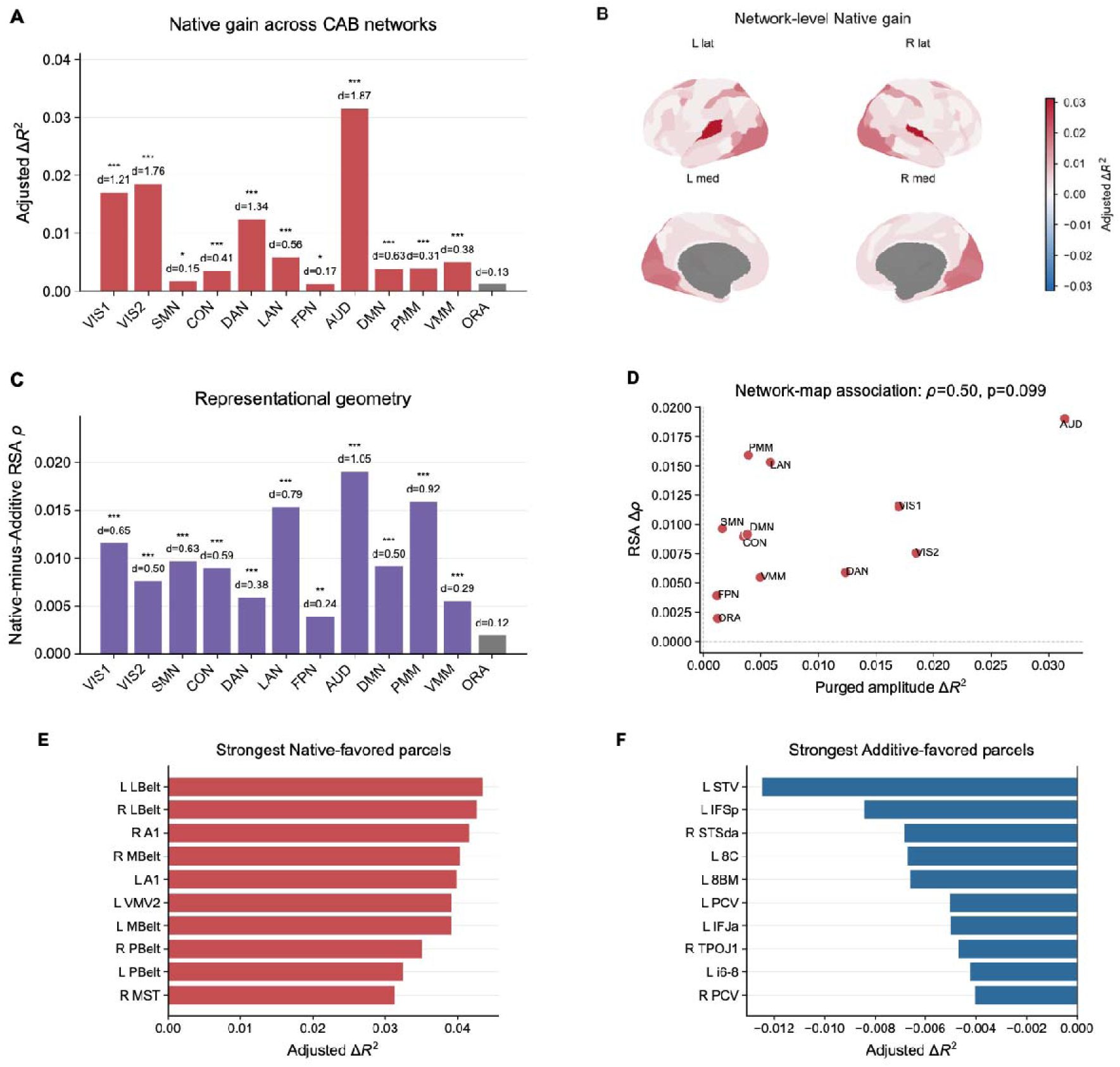
Network hierarchy and complementary representational geometry. (a) Covariate-adjusted NMA in 12 Cole-Anticevic networks; bar height is ΔR² and annotations report two-sided FDR and Cohen’s d. (b) Parcel display of the network estimates using CAB-NP assignments. (c) Native-minus-Additive representational-similarity-analysis effect across networks; bar height is ΔSpearman rho, with positive values indicating that Native feature geometry was closer to brain-pattern geometry than Matched Additive geometry. (d) Association between network amplitude NMA and RSA NMA (Spearman rho=0.50, two- sided p=0.099); zero lines divide effect direction. (e, f) Ten parcels with the largest Native- and Additive- favored adjusted effects. Inference is across 176 participants. AUD, auditory; CON, cingulo-opercular; DAN, dorsal attention; DMN, default; FPN, frontoparietal; LAN, language; ORA, orbito-affective; PMM, posterior multimodal; SMN, somatomotor; VIS1/2, visual 1/2; VMM, ventral multimodal.

A complementary representational-similarity analysis asked whether Native features also better captured the geometry of clip-to-clip brain-pattern relationships. Native-minus-Matched RSA was positive and FDR-significant in 11 of 12 networks (Fig. 3c). The largest RSA differences were in AUD (ΔSpearman rho=0.01906), posterior-multimodal cortex (PMM; 0.01593), language cortex (LAN; 0.01532), and VIS1 (0.01157). Absolute Native and Matched RSA, network-wise RSA differences, run-specific effects, and the amplitude-RSA comparison are reported in Supplementary Fig. S8a-d.

Amplitude NMA and RSA NMA were positively, but not significantly, associated across networks (Spearman rho=0.50, two-sided p=0.099; Fig. 3d). The two analyses therefore converged in overall direction without establishing that amplitude prediction and representational geometry share an identical network ranking. Parcel rankings reinforced this distinction: the largest adjusted Native effects concentrated in auditory and audiovisual sensory regions (Fig. 3e), whereas the largest Additive-favored effects were smaller and distributed across selected frontal, temporal, and parietal parcels (Fig. 3f; Supplementary Fig. S6a, b). Together with the network-size and hemispheric summaries, these controls cover Supplementary Fig. S6a-d. Thus, Native semantics improved both response-amplitude prediction and multivariate structure, but the two signatures were complementary rather than interchangeable.

### Feature replacement reveals bidirectional semantic routing

To identify which semantic axes carried the Native advantage, I performed held-out feature replacement. In each analysis, one Native feature was replaced by its Matched counterpart while all other Native features and the fitted evaluation procedure were retained; replacement loss was defined as R² of the Native model minus R² of the hybrid model (Fig. 4a). Positive loss therefore indicates predictive information specific to the Native version of that feature, whereas negative loss indicates that the Matched feature improved prediction.

**Figure 4.**
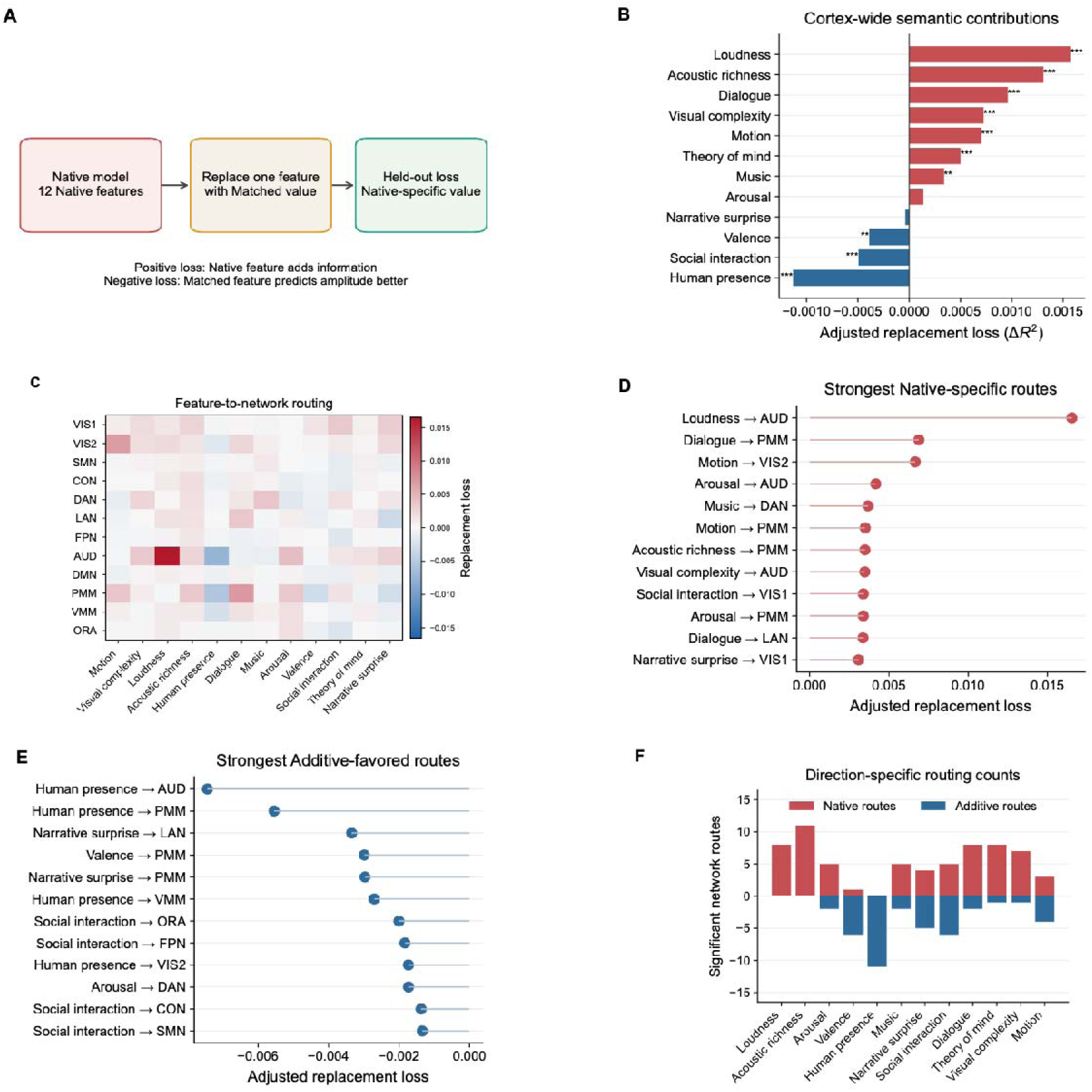
Feature-specific semantic routing accounts for predictive Native gain. (a) Held-out replacement logic. One Native feature was replaced by its Matched counterpart; replacement loss equals R²(Native) minus R²(hybrid), so positive values denote Native-specific predictive information and negative values favor the Matched feature. (b) Cortex-wide participant-adjusted replacement loss for the 12 semantic dimensions; asterisks denote two-sided FDR. (c) Feature-by-network replacement effects across 12 dimensions and 12 CAB-NP networks. (d,e) Largest globally corrected positive and negative routes. (f) Number of globally significant positive and negative routes per dimension. All 144 feature- network tests were corrected together; inference is across 176 participants after adjustment for motion, age, and sex.

At the cortex-wide participant-adjusted level, Acoustic Loudness showed the largest positive replacement loss (0.001573), followed by Acoustic Richness (0.001309), Spoken Dialogue (0.000961), Visual Complexity (0.000726), Visual Motion Energy (0.000701), Theory of Mind (0.000500), and Music Prominence (0.000341); all survived FDR correction (Fig. 4b). In contrast, Human Presence (-0.001132), Social Interaction (-0.000494), and Emotional Valence (- 0.000389) significantly favored their Matched versions. Emotional Arousal and Narrative Surprise were not significant at the cortex-wide level. The mixture of signs rules out a unitary claim that every semantic dimension becomes more brain-predictive when scored jointly.

The feature-by-network matrix revealed substantially richer routing than the cortical averages (Fig. 4c). Of 144 feature-network tests corrected together, 105 were significant: 65 positive Native-specific routes and 40 negative Additive-favored routes. The strongest positive routes included Acoustic Loudness to AUD, Spoken Dialogue to PMM and LAN, Visual Motion Energy to VIS2, Music Prominence to DAN, and Acoustic Richness to PMM (Fig. 4d). The strongest negative routes included Human Presence to AUD and PMM, Narrative Surprise to LAN and PMM, Emotional Valence to PMM, and Social Interaction to several control and association networks (Fig. 4e). Route counts showed that a single feature could be Native-specific in some networks and Additive-favored in others (Fig. 4f).

The full effect, corrected-evidence, and significance matrices, together with the comparison of group-average and participant-adjusted cortical effects, are shown in Supplementary Fig. S9a-d. Parcel-level replacement maps for Visual Motion Energy, Acoustic Loudness, Spoken Dialogue, and Human Presence further demonstrate that each route had a distinct cortical topography (Supplementary Fig. S10a-d). These analyses identify feature-network selectivity, rather than a global semantic-strength shift, as a principal organization of representational non-additivity.

### Native gain is broad across films but gated by content

The overall advantage varied across source films. Among 13 non-repeated sources, nine had positive adjusted Native error gain, and eight were significantly positive after FDR correction (Fig. 5a). The mean adjusted gain across non-repeated films was 0.006238 (two-sided p=0.0002). Northwest Passage, Empire Strikes Back 1, Home Alone 1, and Erin Brockovich 2 showed particularly strong positive effects, whereas Welcome to Bridgeville and Social Network 1 significantly favored Matched Additive. One additional source was positive but non-significant, and two were negative but non-significant, yielding the complete directional outcome counts in Fig. 5b.

**Figure 5.**
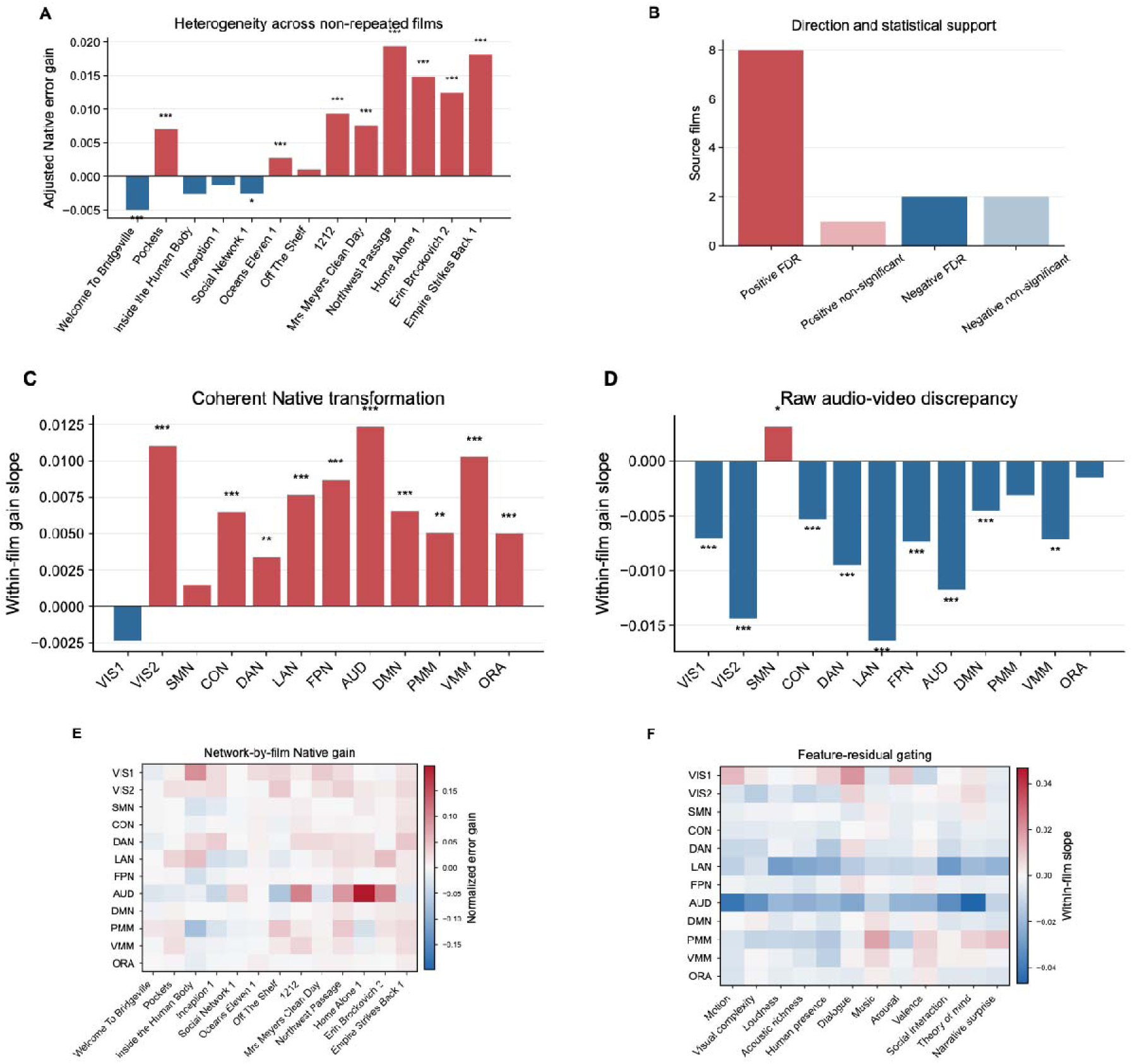
Native gain is broad across films and gated by content. (a) Participant-adjusted normalized error gain in 13 non-repeated source films; red and blue denote Native- and Additive-favored effects, respectively, and asterisks denote FDR across films. (b) Film counts by direction and statistical support. (c,d) Within-film associations of network gain with coherent Native emergence magnitude and raw audio-video discrepancy, estimated from temporally non-overlapping clips with source-film fixed effects. Asterisks denote correction across the 24 prespecified network-mechanism tests. (e) Network-by-film adjusted gain. (f) Feature-residual-by-network association matrix. Color scales encode signed normalized error gain or regression slope. These associations are descriptive of content gating and are not causal estimates.

To test what content properties accompanied this heterogeneity, associations were estimated within source film using temporally non-overlapping clips and source-film fixed effects. Coherent Native emergence magnitude, the magnitude of the joint semantic re-description relative to the Matched representation, was positively associated with Native gain in 10 of 12 networks after correction; VIS1 and somatomotor cortex were not significant (Fig. 5c). By contrast, raw audio-video discrepancy across shared dimensions was negatively associated with gain in VIS1, VIS2, cingulo-opercular, dorsal-attention, language, frontoparietal, auditory, default, and ventral-multimodal networks. Somatomotor cortex showed a smaller positive association, whereas PMM and orbito-affective cortex were not significant (Fig. 5d).

Native gain therefore tracked coherent transformation of multimodal meaning rather than the mere magnitude of disagreement between unimodal descriptions. This interpretation is associational: the analyses identify content gating but do not establish a causal mechanism. The network-by-film matrix showed that each source recruited a different combination of positive and negative network effects (Fig. 5e). Corresponding presentation-block effects, globally corrected network-by-block evidence, all-block estimates, and the four repeated-content presentations are provided in Supplementary Fig. S11a-d.

At the feature level, within-film residual slopes also varied in sign across networks (Fig. 5f). Complete network slopes for emergence magnitude and crossmodal discrepancy, followed by all 144 feature-residual slopes and their globally corrected evidence, are shown in Supplementary Fig. S12a-d. The convergence of film-, network-, and feature-level analyses indicates that the Native advantage was widespread across the stimulus set but conditional on which semantic transformation occurred in which cortical system.

### The effect is temporally robust, but absolute generalization is limited

The Native advantage survived stricter temporal controls. With the primary purge, the group Native-Matched difference was 0.02144; it remained 0.01965 with a 10-s embargo, 0.01963 with a 20-s embargo, and 0.01440 with a 10-s embargo after complete exclusion of the repeated Vimeo clips (Fig. 6a). Participant-adjusted NMA remained positive in every condition (0.007372, 0.006539, 0.006689, and 0.005898, respectively; all corrected p=0.0002), with 85.8- 94.3% of participants showing a positive cortex-wide effect (Fig. 6b). Network-by-condition effects, trajectories of the strongest networks, participant sign consistency, and globally corrected network evidence are shown in Supplementary Fig. S7a-d. Additional checks confirmed the capacity controls, embargo retention, concordant prediction-correlation and R² effects, and stability after excluding participants with mean motion greater than 0.20 (Supplementary Fig. S18a-d).

**Figure 6.**
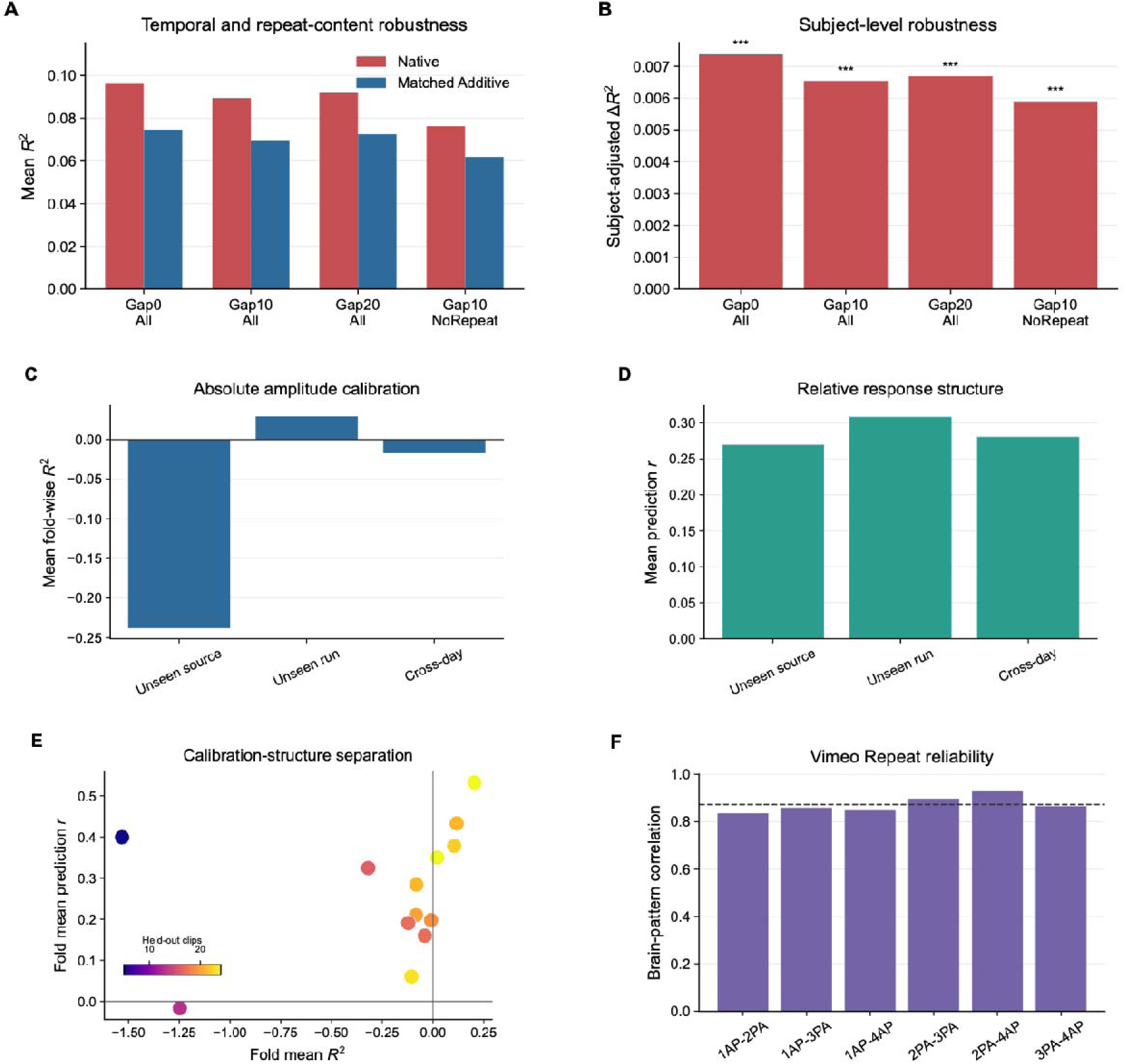
Temporal robustness and the boundary of generalization. (a) Native and Matched Additive group mean R² under the primary purge, 10-s and 20-s embargoes, and a 10-s embargo after repeat exclusion. (b) Participant-adjusted NMA for the same conditions; asterisks denote corrected two-sided bootstrap tests across 176 participants. (c,d) Mean fold-wise R² and prediction correlation for completely unseen source films, unseen runs, and cross-day transfer. Negative R² indicates failure of absolute amplitude calibration, whereas positive r indicates preserved relative response structure. (e) Source-specific R² versus prediction r; point size denotes the number of held-out clips. (f) Pairwise Pearson correlations of flattened group brain patterns during four Vimeo Repeat presentations; the dashed line denotes the six-pair mean.

Generalization beyond the within-film prediction regime exposed a boundary between relative response structure and absolute amplitude calibration. Mean fold-wise R² was -0.2381 for a completely unseen source film, 0.0292 for an unseen run, and -0.0173 for cross-day transfer (Fig. 6c). Mean prediction correlations for the same tests remained positive (r=0.2695, 0.3083, and 0.2804, respectively; Fig. 6d). Thus, models often preserved relative variation in cortical responses even when they failed to reproduce the absolute response scale of a new film or day. This limitation was not spatially uniform. Although cortex-wide mean R² was weak or negative in the most stringent transfer settings, cortical maps revealed localized regions with positive absolute prediction (up to 0.3), indicating that some areas retained meaningful calibration across films, runs, or days (Supplementary Fig. S19a-c).

Source-film folds varied substantially in calibration, response-structure correspondence, and held-out sample size (Fig. 6e). The complete leave-one-source-film-out audit reports mean R², mean r, fold-size dependence, and the calibration-versus-structure plane for every source (Supplementary Fig. S13a-d). Run-specific and bidirectional cross-day transfer, together with the day/run/phase-encoding acquisition map, are shown in Supplementary Fig. S14a-c. Cortical maps further revealed that calibration failure was spatially heterogeneous across unseen-source, unseen-run, and cross-day tests (Supplementary Fig. S19a-c).

The limited absolute transfer was not attributable to an intrinsically unstable repeated stimulus. Group brain patterns evoked by the four Vimeo Repeat presentations were highly reliable, with a mean pairwise pattern correlation of 0.8729 (Fig. 6f). The full repeat correlation matrix, the six individual run pairs, and their minimum/mean/maximum summary are shown in Supplementary Fig. S15a-c.

Finally, individual-difference analyses suggested only a weak trait-like component. Cross-day subject-profile similarity was strongest in VIS1 (r=0.404, q=3.16×10□□), with VIS2 and LAN also surviving FDR, whereas network ICCs were small. Top-1 identification accuracy was 2.27%, above the 0.57% chance level (permutation p=0.0170) but low in absolute terms; top-5 accuracy was 9.09% (Supplementary Fig. S16a-d). Neither prespecified fluid cognition (rho=0.062, q=0.422) nor crystallized cognition (rho=0.105, q=0.344) was associated with cortex-averaged NMA, and all seven secondary cognitive tests were also null (Supplementary Fig. S17a, b). Together, the cognition and construct-distinction controls cover Supplementary Fig. S17a-c. These null incremental associations do not conflict with prior associations involving absolute Gemini explainability because the present estimand is the Native-minus-Additive difference (Supplementary Fig. S17c). Together, the results define a robust within-film representational advantage whose relative structure transfers more readily than its absolute amplitude or participant identity.

## DISCUSSION

This study shows that cortical responses during intact natural audiovisual experience are better predicted by a native joint audiovisual semantic representation than by a dimension-matched account assembled from separate audio-only and silent video descriptions. The central contribution is not a new instance of classical blood oxygen level dependent superadditivity. It is a representational test. I kept the human stimulus fixed and varied the computational description of the same movie clips. This design allowed me to ask whether the cortical response to a natural event preserves information that is aligned with joint audiovisual meaning beyond a matched additive account of unimodal meaning. The result was broad, reproducible, and graded. Native gain appeared across much of the cortex, was strongest in auditory, visual, and dorsal attention systems, and included local regions where the matched additive model performed better. This pattern places the finding within a broader movement from simplified tasks toward naturalistic models of brain function, while also keeping the estimand more specific than general naturalistic prediction^12,14,25–27^.

The methodological motivation is that physical modality removal is a difficult control in continuous movies. Classical audiovisual experiments obtain their force from manipulating auditory and visual inputs, and this logic remains indispensable for causal questions about sensory integration^9,10,28^. Yet in a natural narrative, muting a soundtrack does not only remove auditory energy. It also changes gaze, attention, effort, comprehension, immersion, and the perceived continuity of events^15,16,18,19^. These factors are not nuisances that can be cleanly separated from meaning after the fact. They are part of the viewing state that gives naturalistic paradigms their value. My computational counterfactual approach therefore complements classical modality manipulation. It does not claim that the brain response to the intact movie equals the sum of unmeasured auditory and visual brain responses. Instead, it asks whether alternative semantic descriptions of the same intact experience differ in their ability to predict the same cortical activity. This framing also clarifies why multimodal artificial intelligence is useful here. Deep networks and large multimodal models are not treated as models of the whole human perceptual system. They are used as controlled instruments for generating alternative feature spaces over identical stimuli. This is a narrower and more testable role. In the present case, the Native and Matched Additive spaces shared the same semantic axes and differed in how those axes were estimated. The advantage of NativeAV12 therefore cannot be reduced to a larger feature count, a richer concatenation, or a more flexible regression model. It reflects information in the joint scoring of the intact clip that was not fully recovered by recombining separate unimodal scores. This use of models as explicit representational hypotheses follows a long tradition in encoding analysis and in goal-driven computational neuroscience, while extending that logic to a natural audiovisual semantic setting^24,29–31^.

The cortical distribution of the effect suggests that joint audiovisual semantics are not confined to a small set of classical association regions. The largest gains appeared in auditory cortex, secondary and primary visual cortex, and dorsal attention regions. These systems are often described in sensory or attentional terms, but natural movies place sensory form, action, dialogue, affect, and event structure into a single evolving stream. A representation that scores the intact clip can therefore become more predictive even in regions that are not usually framed as abstract semantic hubs. At the same time, the local additive favored parcels are scientifically useful. They show that the analysis did not simply reward Native scores everywhere. Instead, cortex contained a dominant Native advantage with meaningful local exceptions. This fits with evidence that natural meaning is distributed across large cortical maps and that semantic representations can be partially shared across input formats while still retaining modality- and region-dependent structure^26,32,33^. The representational similarity analysis sharpened this interpretation. Native features improved not only response amplitude prediction, but also the geometry of clip-to-clip response relationships across networks^26,29,34^. This matters because a model could in principle predict mean response levels while failing to capture how different events are organized relative to one another. The RSA results indicate that Native scores carried information about relational structure as well as amplitude. Yet amplitude gain and RSA gain were only moderately aligned across networks and their map association did not reach conventional significance. I therefore interpret the two analyses as complementary views of representational fit, not as redundant measures of a single hidden variable. This distinction is important because naturalistic brain responses can vary in baseline amplitude, reliability, and relational structure, and each property can constrain a different aspect of model validity^29,34^.

The feature replacement results provide the main organizational principle of the study. Native gain was not a global label that applied equally to all dimensions. It was routed through specific feature and network combinations. Acoustic loudness, acoustic richness, spoken dialogue, visual complexity, visual motion energy, theory of mind, and music prominence carried positive Native specific information at the cortical level. Human presence, social interaction, and emotional valence showed significant matched additive advantages. At the network level, the same semantic axis could change sign across systems. This bidirectional routing argues against a simple fusion narrative in which joint audiovisual scoring is always better. It instead suggests that natural audiovisual semantics are decomposed into multiple cortical uses. Some systems benefit from the integrated interpretation of the intact stream, whereas others are better predicted by modality-separated estimates that preserve simpler or more diagnostic cues^26,32,33^. The source film and content gating analyses add a second principle. Native gain was broad across films, but it was not content invariant. It increased when the intact audiovisual clip produced coherent semantic emergence relative to the matched additive description. It tended to decrease when audio-only and video-only scores diverged in their raw shared dimensions. This dissociation is conceptually important. It means that a large discrepancy between modalities is not sufficient to create a Native advantage. The critical condition appears to be whether the intact audiovisual event supports a useful joint reinterpretation of meaning. This interpretation is consistent with work showing that continuous narratives are organized by event structure and by temporally extended integration windows, but my evidence remains associational^19,35–37^. The present data show content gating of prediction gain rather than a causal mechanism of perceptual binding.

The robustness analyses define the range over which the main claim is defensible. Native advantage survived longer temporal embargoes and the complete removal of repeated Vimeo clips. This makes it unlikely that the result was driven only by adjacent overlapping clips, repeated content, or a leakage artifact of the cross-validation design. At the same time, the generalization analyses exposed a clear boundary. Prediction correlations remained positive for unseen source films, unseen runs, and cross-day transfer, whereas absolute R squared was often negative for unseen sources and across days. I read this as a distinction between preserving relative response structure and calibrating absolute response amplitude. The former was more stable than the latter. This boundary is not a weakness to hide. It prevents overclaiming and places the result in the correct methodological category. The study provides a strong within-film representational comparison under strict purging, not a fully calibrated universal predictor of arbitrary movie responses^7,29,38,39^. The individual difference and cognition analyses further constrain interpretation. Cross-day fingerprinting showed evidence for a weak trait-like component, especially in visual networks, but absolute identification accuracy remained low^7^. Fluid and crystallized cognition were not significantly associated with cortex-averaged NMA. These null results should not be taken as evidence that movie semantics lack behavioral relevance. They show that the incremental Native minus Additive estimand did not explain the selected cognitive measures in this sample. This differs from my prior work on absolute Gemini explainability, which asked how much cortical activity could be predicted by an AI semantic model rather than how much additional prediction was gained by native joint audiovisual scoring over a matched additive baseline^24^. Treating these as distinct neural estimands is essential for avoiding apparent contradictions across studies.

Several limitations follow directly from this framing. First, participants did not view human auditory-only or visual-only versions of the stimuli. The study therefore cannot establish classical neural superadditivity, nor can it estimate the causal effect of removing one sensory channel from perception^9,28^. Second, the semantic axes were defined through Gemini and through a single operational baseline. Other multimodal models, prompts, scoring schemes, or matched additive constructions may reveal different forms of nonadditivity. Third, the stimulus set contained a limited number of source films, and generalization of absolute amplitude to unseen sources was weak. Fourth, the group-derived prediction operators were applied to participants to support subject-level inference, but this is not the same as an independently trained individual predictive biomarker. These constraints are not incidental. They delimit the contribution. My study shows that intact natural audiovisual cortical responses contain feature- and network-specific predictive information aligned with native joint semantic descriptions. It also shows that this information is content-gated and only partly generalizes in absolute amplitude. By keeping the natural viewing experience intact and moving the manipulation into the computational representation, the work offers a tractable framework for studying multisensory semantic organization without disrupting the experience that gives naturalistic cognition its structure^12,31,40^.

## METHODS

### Participants and ethics

This study used only publicly released Human Connectome Project Young Adult data. The original data collection was approved by the Washington University Institutional Review Board, and all participants gave written informed consent. I did not acquire new human data for this study.

I analysed the 176 HCP Young Adult participants who completed all four 7 T movie-watching fMRI runs. The sample contained 70 male and 106 female participants between 22 and 35 years of age. Imaging and behavioural files were downloaded from the HCP distribution^41^.

### MRI acquisition, preprocessing, and parcel time series

The analysis used the HCP structural MRI and 7 T movie watching fMRI data. The relevant fMRI series were acquired with gradient echo EPI at TR = 1000 ms, TE = 22.2 ms, and 1.6 mm isotropic spatial resolution. The structural reference images were 3 T T1-weighted MPRAGE scans with TR = 2400 ms, TE = 2.14 ms, and 0.7 mm isotropic spatial resolution^42^.

I used the HCP minimally preprocessed surface data^43^. In brief, the HCP pipeline includes nonlinear registration of the T1-weighted image to MNI space, cortical surface reconstruction, multimodal surface registration, correction of EPI distortions, motion correction, registration to the structural image, projection to the 32k fs LR CIFTI surface space, and ICA FIX denoising^44 45 46 47 48^.

Movie time series were summarized in the HCP MMP1.0 cortical parcellation. The atlas contains 360 cortical parcels, 180 in each hemisphere, in the same 32k fs LR surface space as the preprocessed CIFTI data^49^. For each participant and run, I extracted parcel mean time series and retained the HCP framewise motion estimates. The mean relative RMS displacement across the four movie runs was used as a subject-level motion covariate in all subject-level inferential analyses together with age and sex.

### Movie stimuli, clip generation, and brain patterns

Participants viewed four HCP 7 T movie runs. The runs contained Creative Commons excerpts and professionally edited Hollywood excerpts separated by 20 s fixation intervals. Participants were instructed to watch naturally and made no behavioural responses during movie viewing^50^.

I reconstructed the stimulus timing from HCP movie metadata and generated overlapping clips from the non rest portions of the four runs. For each continuous movie block, clips were targeted at 20 s duration, allowed to shorten to a minimum of 16 s near block boundaries, and advanced in 10 s steps, corresponding to 50% temporal overlap. A 5 s terminal buffer was reserved so that the later haemodynamic response window remained within the movie block. Clips were exported with FFmpeg at 24 frames per second with synchronized audio. This procedure produced 293 raw audiovisual clips.

The stimulus hierarchy was then verified at the source film level. Each clip was mapped to run, presentation block, source title, stimulus family, day, session, phase encoding direction, and source relative time. Clips from a movie1AP interval with visually confounding subtitles were excluded before modelling. The final encoding sample contained 270 clean clips in 17 presentation blocks, spanning 14 source identities. Thirteen identities were non-repeated source films, and one common Vimeo source was repeated in four presentations.

For Gemini scoring controls, I also generated unimodal derivatives from the audiovisual clips. Video-only clips preserved the video stream and removed audio. Audio-only clips were represented as black screen videos with the original audio stream and the same spatial resolution as the parent clip, allowing the multimodal API to process the audio without informative visual content.

For each subject, I aligned the 270 clean clips to the parcel time series with a 5 s haemodynamic lag. Within each run, parcel time series were z-scored using non-rest frames only. I then subtracted the frame-wise cortical mean across parcels to reduce global fluctuations. For each clip, the lagged response window was averaged across TRs, yielding a 360- dimensional response vector. Group response patterns were computed as a 5% trimmed mean across subjects for each clip and parcel, while the individual clip patterns were retained for subject-level inference.

### Gemini semantic scoring and feature spaces

Each clip was scored with a custom Gemini 3.1 Pro (Google DeepMind) annotation pipeline. The API received one video file and a fixed zero-shot instruction that requested continuous scores from 0.00 to 1.00, rounded to two decimal places, together with a brief reason for each score. Responses were constrained to a JSON schema and saved as both JSON reasoning files and score CSV tables. Failed or missing clips were retried, and the final CSV tables were generated from the stored JSON outputs.

Three score tables were used. The NativeAV table was obtained from intact audiovisual clips. The Audio table was obtained from black screen audio clips with the original soundtrack. In that condition, the prompt instructed Gemini to use acoustic evidence. The Video table was obtained from silent video clips. The 12 dimensions were designed to include visual-specific dimensions, auditory-specific dimensions, and higher-level shared semantic dimensions that could plausibly be estimated from either modality.

I compared seven prespecified feature spaces. Audio9 contained the nine dimensions available from the audio-conditioned score table after excluding visual-specific dimensions. Video8 contained the eight dimensions available from the silent-video score table after excluding auditory-specific dimensions. NativeAV12 used the 12 intact audiovisual scores. MatchedAdditive12 was the primary control and had the same 12 dimensions as NativeAV12. Its construction followed the formula below, where Mj is the MatchedAdditive12 value for feature j, Aj is the audio-conditioned score, and Vj is the silent-video score.

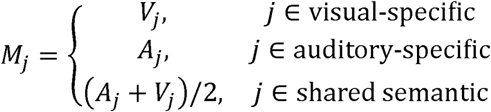

The visual-specific set comprised Visual Motion Energy, Visual Complexity, and Human Presence, so these dimensions used silent-video scores. The auditory-specific set comprised Acoustic Loudness, Acoustic Richness, Spoken Dialogue, and Music Prominence, so these dimensions used audio-conditioned scores. The shared semantic set comprised Emotional Arousal, Emotional Valence, Social Interaction, Theory of Mind, and Narrative Surprise, so these dimensions were averaged across audio-conditioned and silent-video scores. This design made NativeAV12 and MatchedAdditive12 identical in dimensionality and feature names while differing in whether each semantic axis was estimated from intact audiovisual context or from a matched unimodal semantic counterfactual.

Three additional controls tested whether NativeAV12 benefited from dimensionality or concatenation rather than joint scoring. LateConcat17 concatenated the modality-specific Audio9 and Video8 features. ConcatPCA12 applied train-only PCA to the 17 concatenated features and retained 12 components. LegacyConcat24 concatenated the full 12 audio and 12 video score vectors. All feature spaces were aligned to the same clean clip IDs.

### Purged ridge encoding and statistical inference

The primary estimand was Native minus Matched Additive predictive performance. For each held-out clip, I removed from the training set every clip that had the same source title and overlapping source relative time. This purge removed the held-out clip itself, temporally overlapping neighbours generated by the 50% sliding window, and aligned repeated Vimeo content when the same source appeared in another run.

Ridge models were fit separately for each cortical parcel, but all parcels were predicted with the same fold-specific linear operator. Features were standardized using training clips only. Responses were centred using training clips only. For PCA controls, the PCA rotation was also estimated only from the training features. Ridge penalties were selected from logspace(-3, 5, 17). For every source identity, the complete source was excluded from the inner lambda selection procedure, and the selected lambda was then used to predict clips from that source with all non-purged training observations. Prediction was quantified as out-of-sample R-squared and Pearson prediction correlation across clips.

I first estimated all seven feature spaces on the group-averaged clip patterns. I then fixed the NativeAV12 and MatchedAdditive12 prediction operators from the group analysis and applied them to each subject independently. This produced subject-by-parcel maps of Native R squared, Matched Additive R squared, prediction correlations, and Native minus Additive R squared. I refer to the primary difference as NMA.

Subject-level inference used the 176 subject NMA maps. For parcel-level, network level, film level, feature replacement, robustness, RSA, and cognition analyses, I adjusted for mean head motion, age, and sex. The intercept of the adjusted model was tested with a Rademacher wild bootstrap^51^ applied to residuals from the reduced covariate model. The t statistic was the intercept estimate divided by its standard error from the covariate-adjusted linear model. Unless otherwise stated, 5000 bootstrap samples were used. I report two-sided p values, Benjamini-Hochberg FDR-corrected q values^52^, and max statistic family-wise error p values where the analysis defined a family of spatial or feature tests.

Cortical network summaries used the CAB NP assignment of HCP MMP parcels into 12 networks^49,53^. Network effects were computed by averaging parcel-wise subject metrics within each network before covariate-adjusted inference. Cohen d was computed from the unadjusted subject-level effect distribution and is used as a standardized descriptive effect size.

### Representational similarity and feature replacement

I tested representational geometry independently of the encoding amplitude analysis. Within each run, I selected non-overlapping clips by advancing through source time and retaining the next clip only when it began after the previous retained clip ended. For each network and subject, I computed a brain representational dissimilarity vector as pairwise correlation distance among clip response patterns. NativeAV12 and MatchedAdditive12 feature dissimilarity vectors were computed from z-scored features for the same selected clips. Spearman correlations between brain and model dissimilarities produced Native and Matched RSA values, and their difference defined RSA NMA. Positive RSA NMA indicates that Native feature geometry was closer to brain-pattern geometry than Matched Additive geometry. Network-level RSA inference used the same covariates and bootstrap framework as the amplitude analysis.

To identify which semantic dimensions carried Native specific information, I created 12 hybrid models. In each hybrid model, one NativeAV12 feature was replaced by the corresponding MatchedAdditive12 feature while all other Native features were left unchanged. Each hybrid model received its own source-excluded lambda selection and the same content-aware purge used in the primary analysis. Replacement loss was defined as NativeAV12 R squared minus hybrid R squared. Positive loss indicates information carried by the Native feature that was not recovered by its matched additive counterpart. Feature replacement inference was performed at cortex-wide and CAB network levels. The 144 feature-by-network tests were corrected as a single family.

### Source film, content gating, robustness, and generalization

To test whether Native gain varied across movie content, I summarized purged predictions within each clean presentation block. For each subject and block, error gain was defined as the Matched Additive squared error minus the NativeAV12 squared error, divided by response variance estimated from clips outside the same source identity. This scaling prevented the test source from defining its own normalization. Inference was across subjects for each block and for each network by block combination.

Content gating analyses focused on temporally non-overlapping clips from the 13 non-repeated source films. I fitted within-film models with source film fixed effects. Two mechanism predictors were tested. Native semantic emergence was the root mean squared residual between NativeAV12 features and MatchedAdditive12 features after feature standardization. Crossmodal discrepancy was the root mean squared difference between standardized audio-only and video-only scores over the shared semantic dimensions. I also tested each Native minus Matched feature residual separately. Slopes were estimated within-subject and network and then tested across subjects with the covariate-adjusted bootstrap procedure.

Robustness analyses repeated the NativeAV12 and MatchedAdditive12 comparison under four prespecified conditions. The primary condition used no additional embargo and retained all clean clips. Additional analyses imposed 10 s and 20 s temporal embargoes around the held-out source relative interval. A fourth analysis used a 10 s embargo after excluding all common repeat clips. Lambda values remained the source cross-fitted values from the primary group analysis.

Generalization analyses evaluated NativeAV12 absolute prediction outside the primary Native versus Additive contrast. The common Vimeo repeat was excluded. I fit leave one source film out models, leave-one-run-out models, and cross-day transfer models. These analyses were summarized with both R squared, which measures absolute amplitude calibration, and Pearson prediction correlation, which measures preservation of relative response structure.

The repeated Vimeo content was used to estimate group response reliability. Repeat clips were aligned by source relative start time across the four presentations, and reliability was computed as the correlation between flattened group brain pattern matrices for each pair of repeat presentations.

### Individual stability and cognition

To test whether NMA contained a subject-stable component, I used the source block by network gain estimates. Within each block and network, values were z-scored across subjects to remove block scale. Day 1 and day 2 network profiles were then averaged separately. Cross-day reliability was summarized by network Pearson correlations and by subject fingerprinting, where each day 1 profile was matched to the most similar day 2 profile. A 10000 permutation null distribution was used for top 1 identification accuracy.

Cognition analyses were restricted to prespecified HCP measures. For each subject, I computed a cross-fitted top decile NMA summary by selecting the top 10% of parcels from the mean NMA map of all other subjects and then averaging the held subject values in those parcels. The two primary outcomes were age-adjusted fluid cognition and age-adjusted crystallized cognition. Secondary outcomes were PMAT24 correct responses, episodic memory, working memory, cognitive flexibility, inhibitory control, theory of mind task performance, and story comprehension accuracy. Associations were partial Spearman correlations controlling for motion, age, and sex, with FDR correction within the primary and secondary families.

## Data and code availability

The MRI data used in this study are available from the Human Connectome Project database. https://www.humanconnectome.org/

The custom MATLAB and Python scripts used for stimulus preparation, Gemini score alignment, encoding analyses, inference, robustness analyses, and figure generation are organized with the manuscript files and uploaded as supplement files.

Software and resources used in this study include Gemini, Google AI Studio, FFmpeg, CIFTI, GIFTI, HCP Workbench, MATLAB, Python, NumPy, pandas, SciPy, Matplotlib, nilearn compatible surface resources.

## ACKNOWLEDGEMENTS

I would like to thank Dr. John Gore and Dr. Zhaohua Ding for their valuable advice and inspiring discussions that have broadly influenced my research.

Imaging data were provided by the Human Connectome Project, WU-Minn Consortium (Principal Investigators: David Van Essen and Kamil Ugurbil; 1U54MH091657), funded by the 16 NIH Institutes and Centers that support the NIH Blueprint for Neuroscience Research; and by the McDonnell Center for Systems Neuroscience at Washington University.

## AUTHOR CONTRIBUTIONS

M.L.: Writing, Visualization, Validation, Software, Methodology, Investigation, Conceptualization.

