## Supplement Figures for "Computational Counterfactuals Reveal Non-Additive Audiovisual Semantics in Natural Movie Responses"

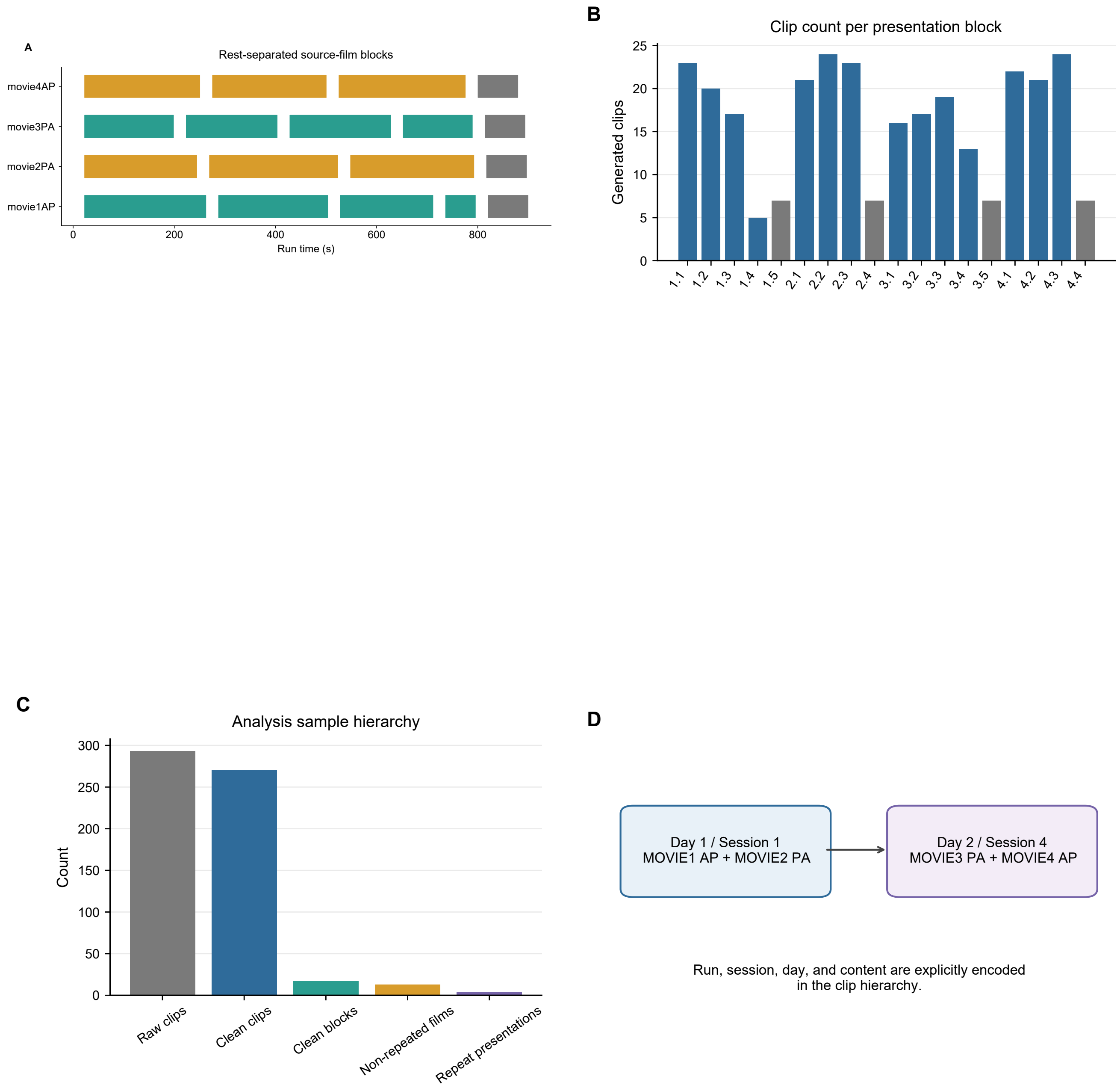


**Supplementary Figure 1 | Stimulus provenance and acquisition hierarchy.** (A) Rest-separated source-film blocks within the four movie runs. Gold marks ordinary source-film excerpts, teal marks Creative Commons material, and grey marks the common Vimeo Repeat; horizontal position is run time in seconds. (B) Number of generated 20-s clips in each presentation block; grey bars identify repeat presentations and blue bars all other blocks. (C) Sample attrition and hierarchy from 293 raw clips to 270 clean clips, 17 clean presentation blocks, 13 non-repeated source films, and four repeat presentations. (D) Mapping of movie runs to scanning day, session, and AP/PA phase encoding; the arrow denotes the cross-day acquisition sequence.

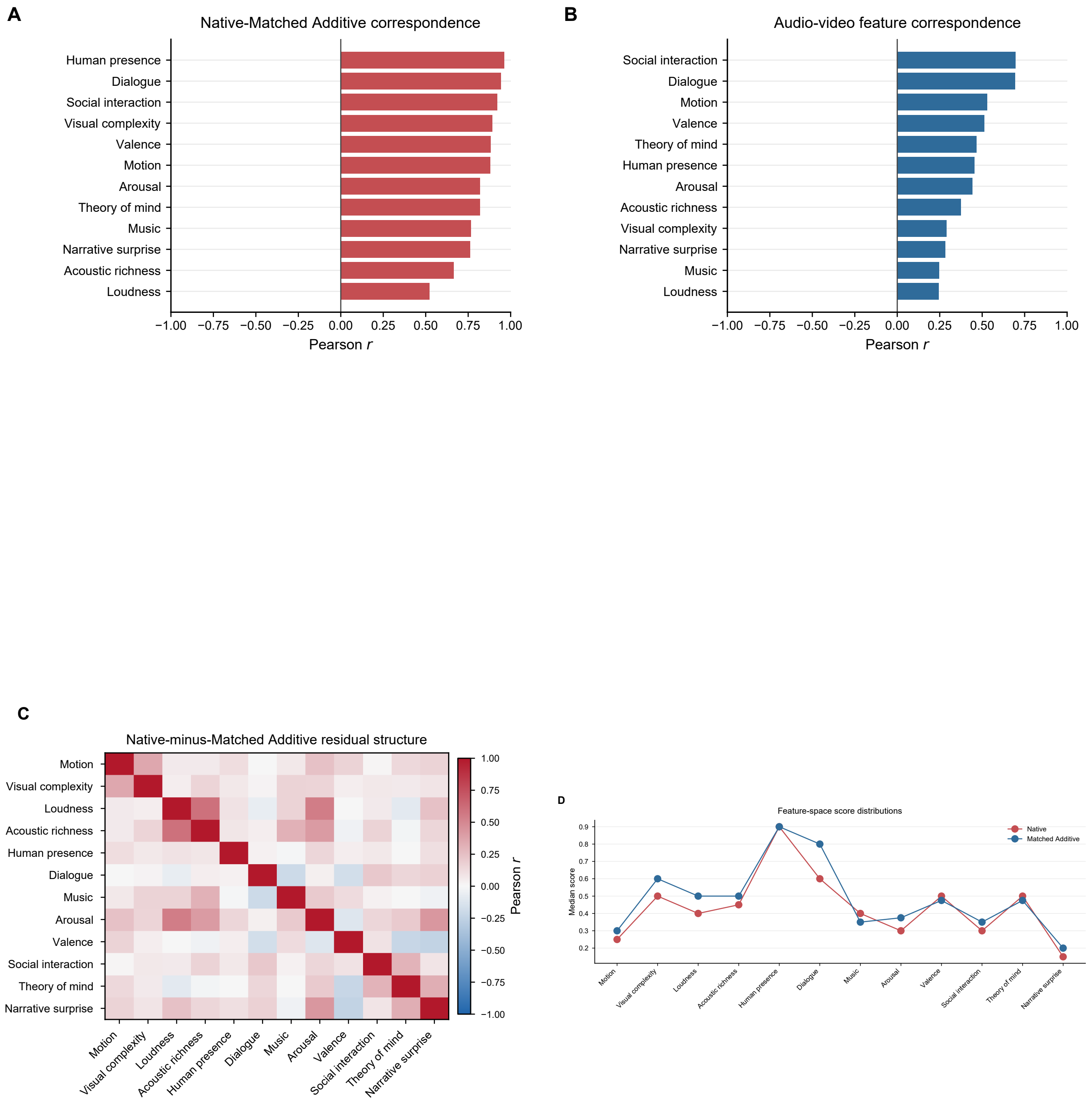


**Supplementary Figure 2 | Structure of Native, audio, video, and Matched Additive semantic spaces.** (A) Pearson correlation between Native and Matched Additive scores for each semantic dimension; the vertical line denotes zero. (B) Pearson correlation between audio-only and video-only scores for the same dimensions. (C) Correlation matrix of Native-minus-Matched Additive feature residuals; the diverging color bar gives Pearson r, with white denoting zero. (D) Median feature scores for Native and Matched Additive spaces; connected red and blue points facilitate dimension-wise comparison.

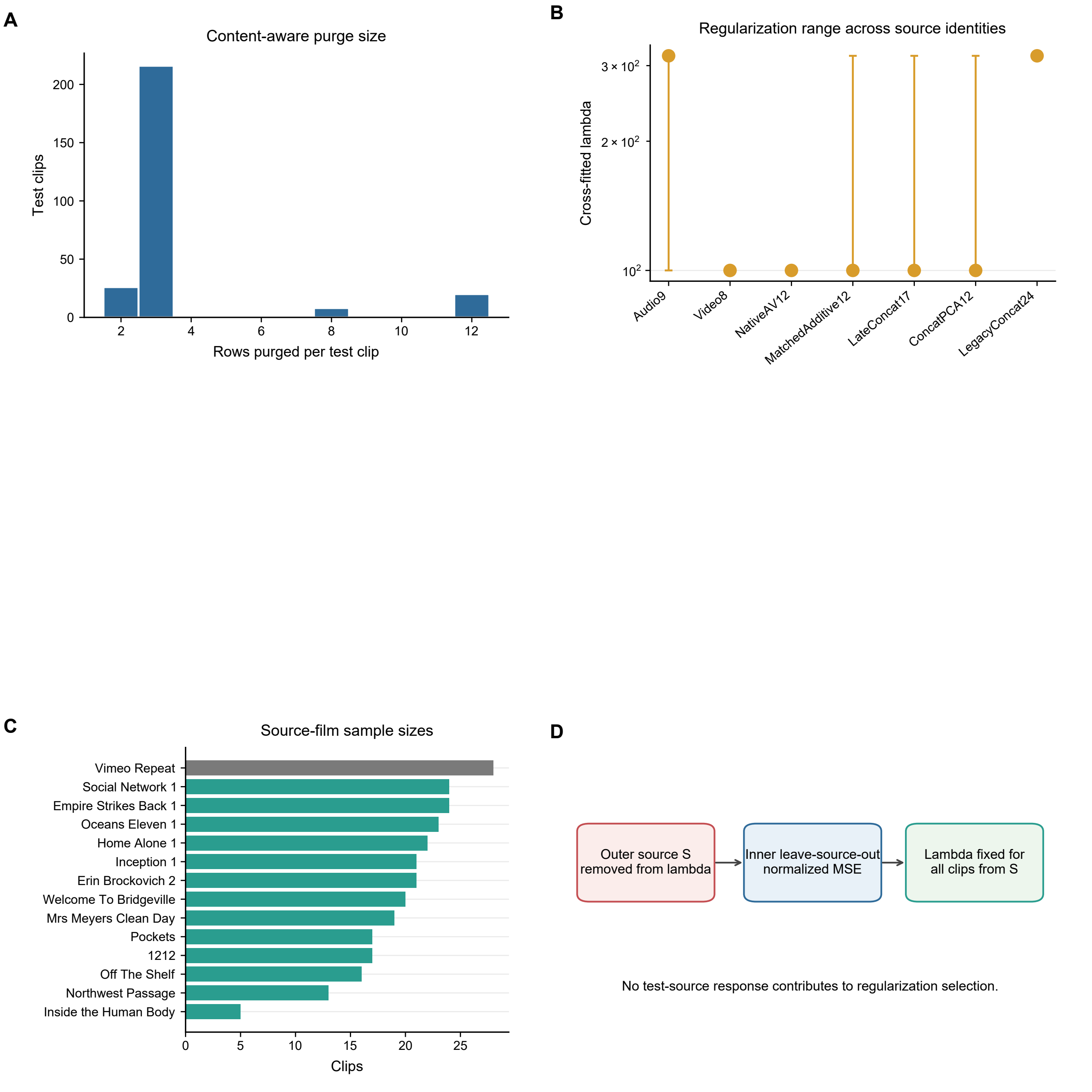


**Supplementary Figure 3 | Purging and regularization diagnostics.** (A) Distribution of the number of response rows removed for each held-out clip by content-aware purging. (B) Median cross-fitted ridge lambda for each model; whiskers span the observed minimum to maximum and the ordinate is logarithmic. (C) Number of clean clips contributed by each source film; the common repeat is grey and non-repeated films are teal. (D) Nested source-excluded lambda-selection scheme. Arrows indicate the order of operations; no response from the outer held-out source enters regularization selection.

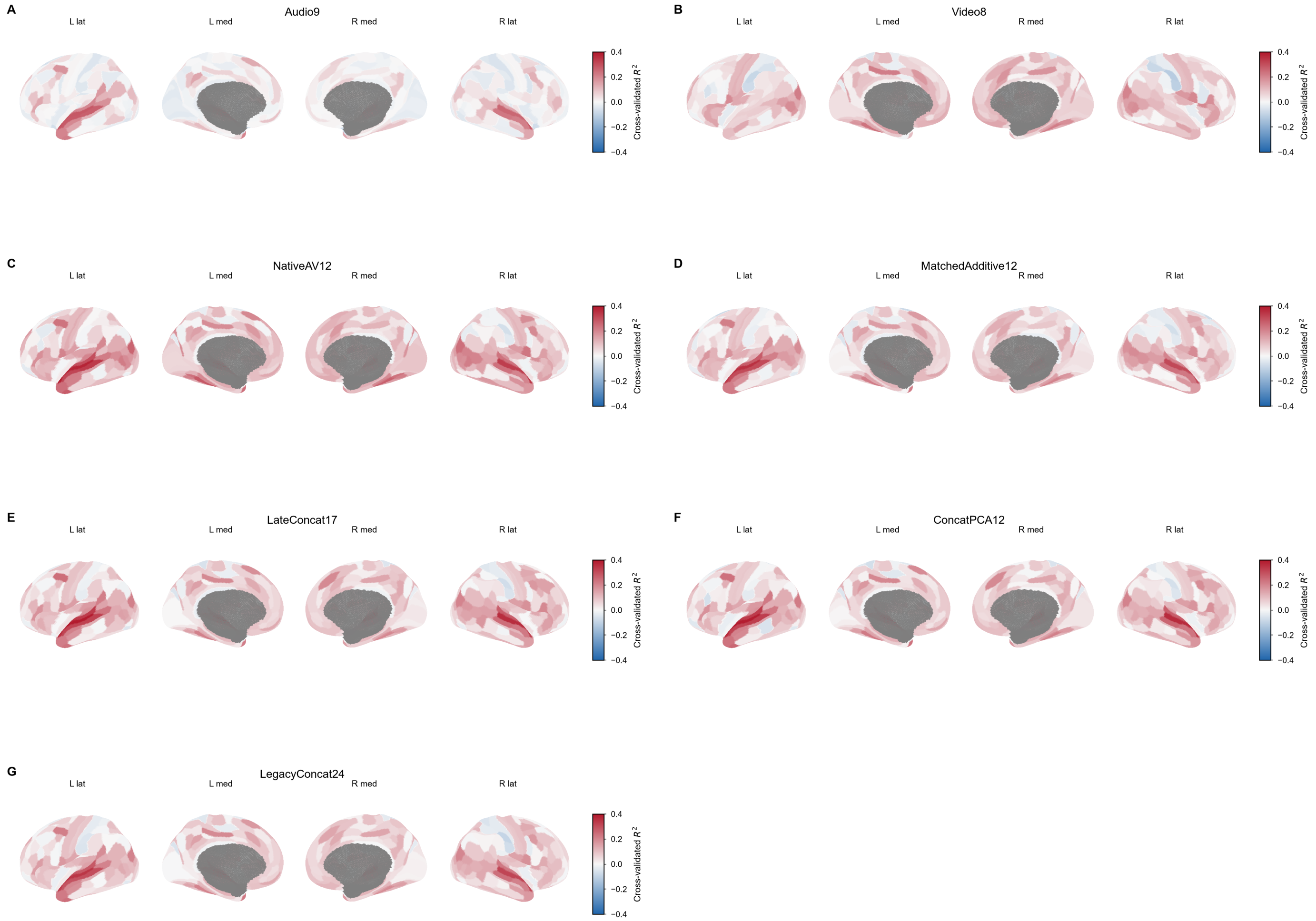


**Supplementary Figure 4 | Cortical surface maps of cross-validated $R^2$ for all seven encoding models.** (A) Audio-only Gemini, nine dimensions. (B) Video-only Gemini, eight dimensions. (C) intact Native audiovisual Gemini, 12 dimensions. (D) dimension-matched modality-wise additive model, 12 dimensions. (E) late concatenation, 17 dimensions. (F) PCA-reduced concatenation, 12 dimensions. (G) legacy concatenation, 24 dimensions. Every panel shows left lateral, left medial, right medial, and right lateral views and uses the identical -0.40 to 0.40 color scale; blue denotes negative out-of-sample $R^2$, white zero, and red positive $R^2$.

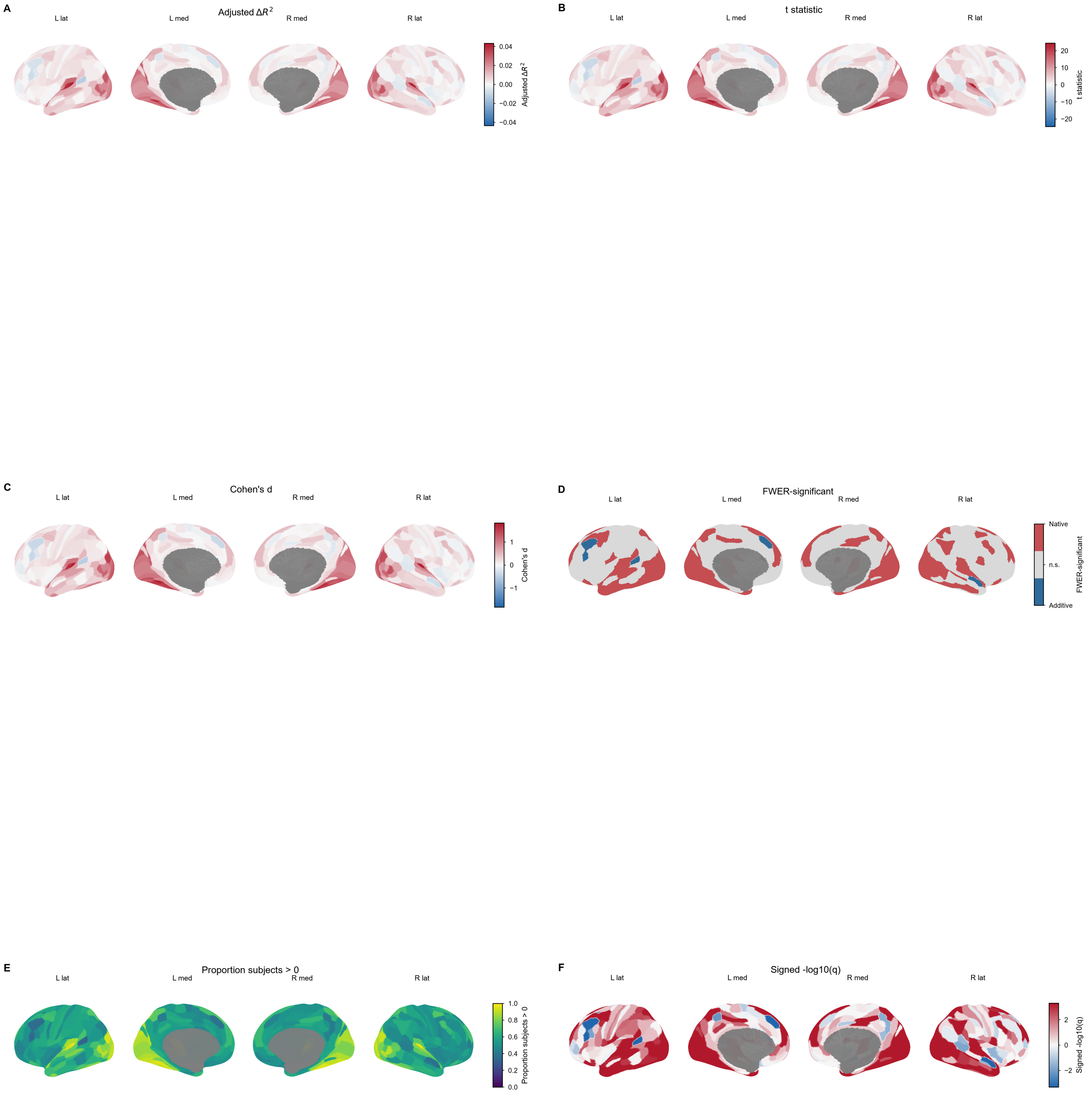


**Supplementary Figure 5 | Complete parcel-level Native-minus-Additive inference.** (A) Covariate-adjusted mean $\Delta R^2$ across 360 MMP parcels. (B) parcel-wise t statistic. (C) Cohen's d, the standardized effect size across 176 subjects. (D) direction of parcels surviving max-statistic family-wise error correction; red and blue denote Native- and Additive-favored parcels and zero-valued grey cortex is not significant. (E) proportion of subjects with positive NMA, shown on a 0-1 sequential scale. (F) signed −log10(two-sided FDR q), where sign follows adjusted $\Delta R^2$. Surface color bars report the quantity named in each panel.

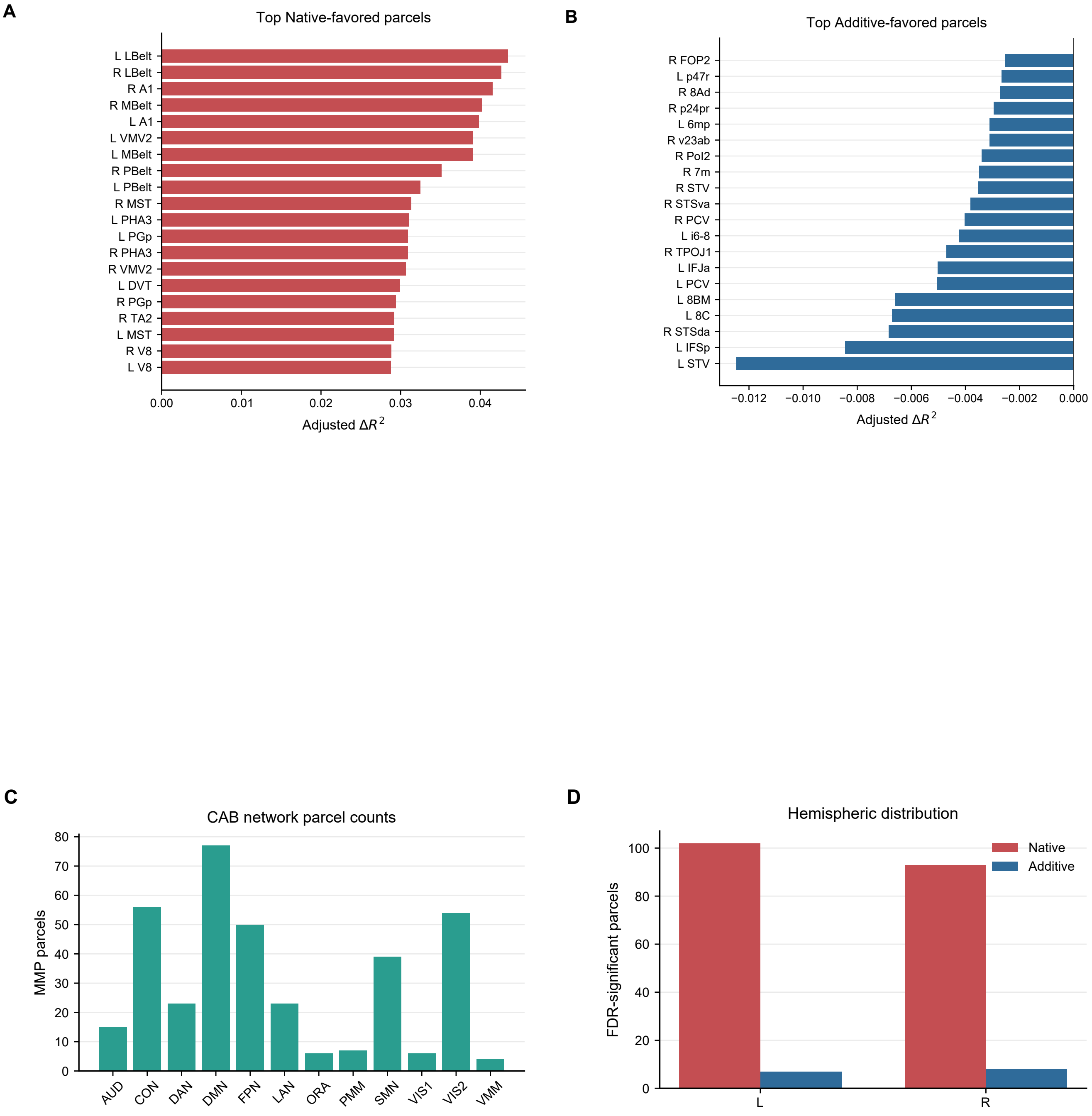


**Supplementary Figure 6 | Ranked regional effects and anatomical balance.** (A) Twenty largest Native-favored adjusted parcel effects; bar length is adjusted ΔR² and the zero line denotes equality. (B) Twenty largest Additive-favored effects on the same estimand. (C) number of MMP parcels assigned to each Cole-Anticevic network, documenting unequal network size. (D) counts of FDR-significant Native- and Additive-favored parcels in the left and right hemispheres; red and blue encode effect direction.

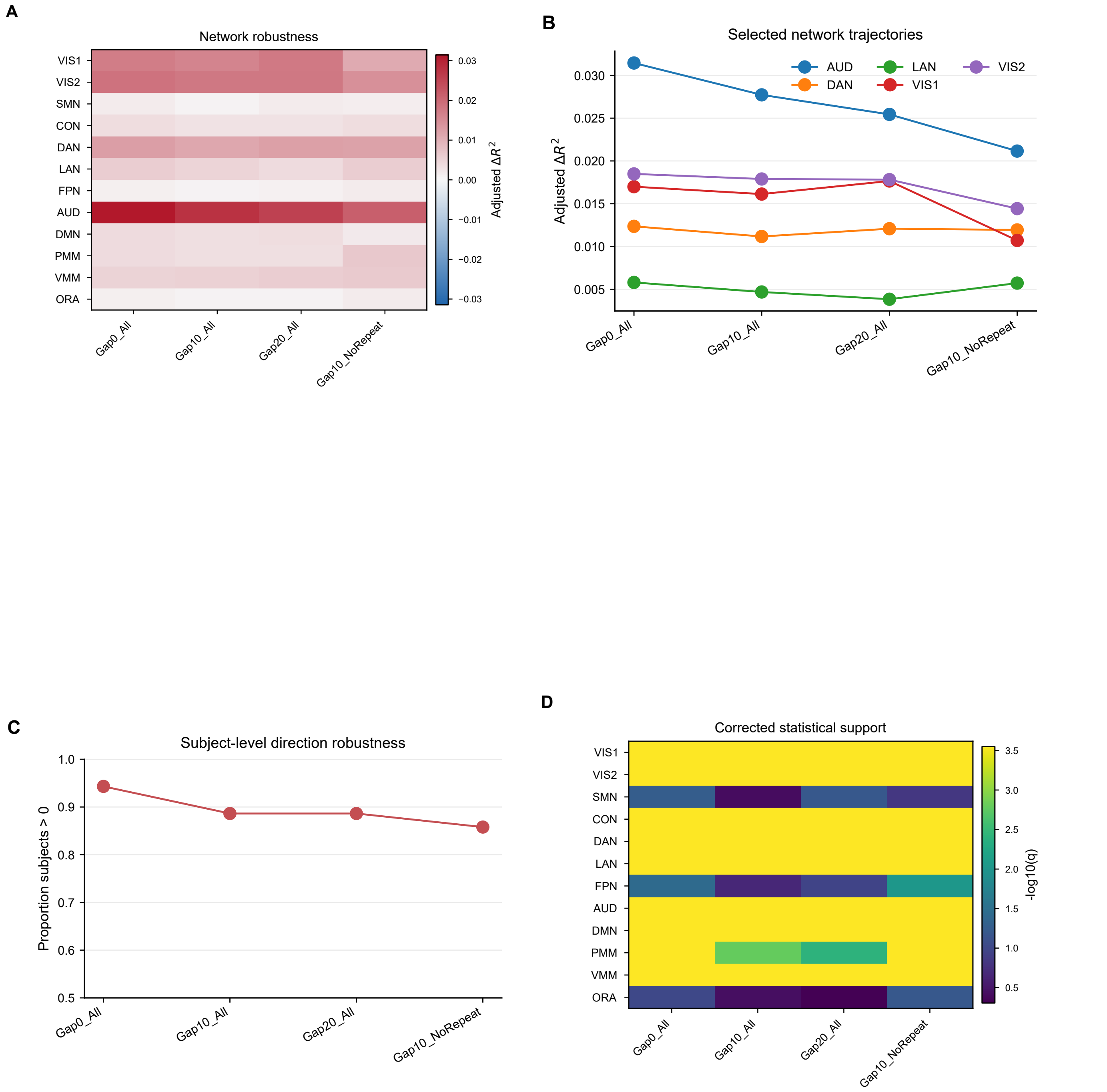


**Supplementary Figure 7 | Network robustness across temporal embargo and repeat-exclusion conditions.** (A) CAB-network by robustness-condition matrix of covariate-adjusted ΔR²; the diverging color bar is the effect-size scale and white denotes zero. (B) effect trajectories for AUD, VIS1, VIS2, DAN, and LAN across the same conditions; points are adjusted network effects. (C) proportion of subjects with cortex-averaged NMA greater than zero; 0.5 is the direction-chance reference. (D) −log10 globally corrected q for all network-condition tests; larger values indicate stronger evidence and q=0.05 corresponds to −log10(q)=1.30.

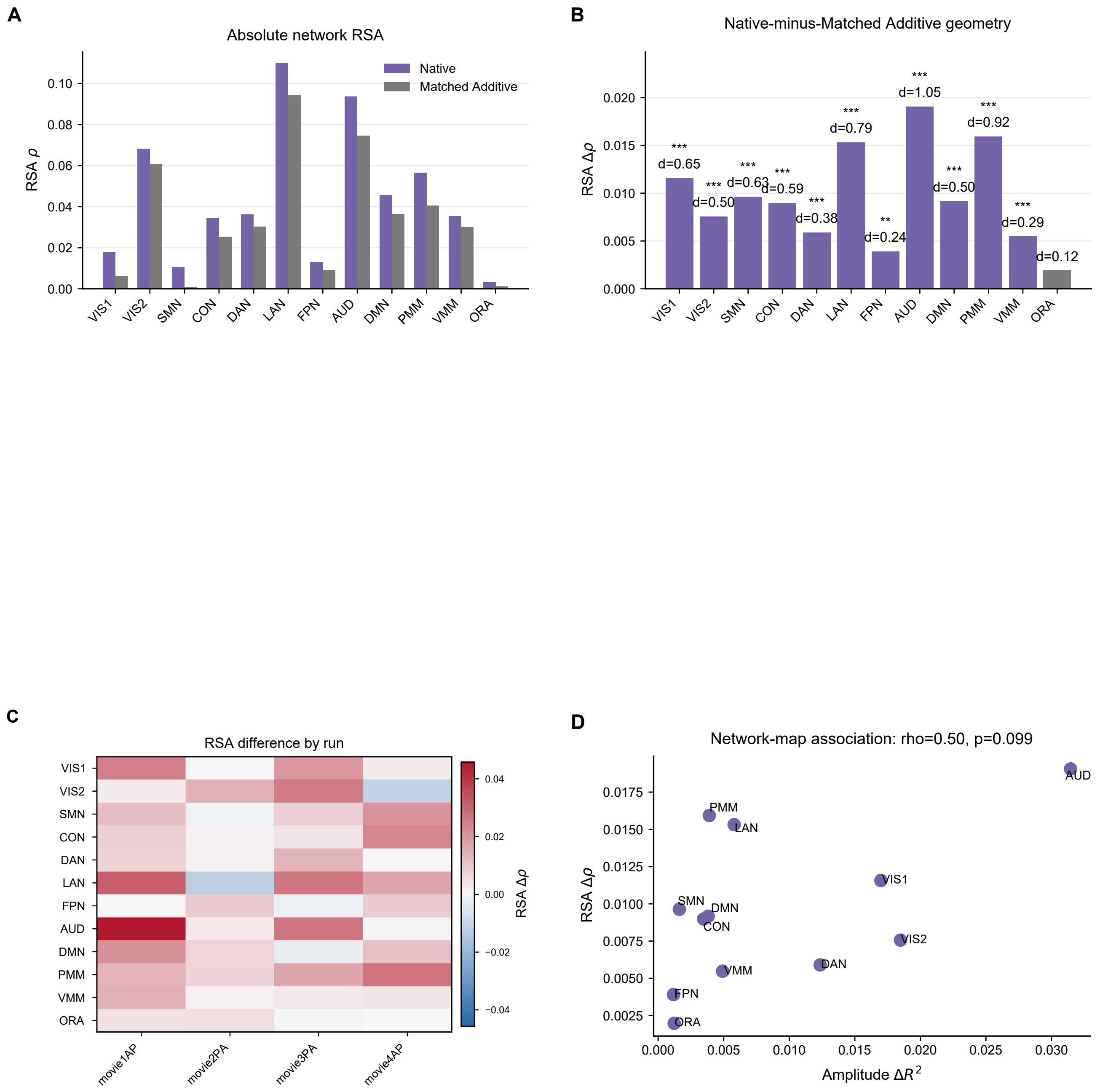


**Supplementary Figure 8 | Complete representational-similarity results.** (A) Absolute Native and Matched Additive network RSA, expressed as Spearman rho. (B) Native-minus-Matched Additive RSA effect; bar height is Δrho, asterisks denote two-sided FDR, and Cohen's d is printed beneath significant symbols. (C) network Δrho separately for each movie run; the diverging color bar denotes direction and magnitude. (D) association between network amplitude $\Delta R^2$ and RSA Δrho; zero lines divide concordant and discordant quadrants, network abbreviations label points, and the title reports the positive but non-significant Spearman map correlation and two-sided p value.

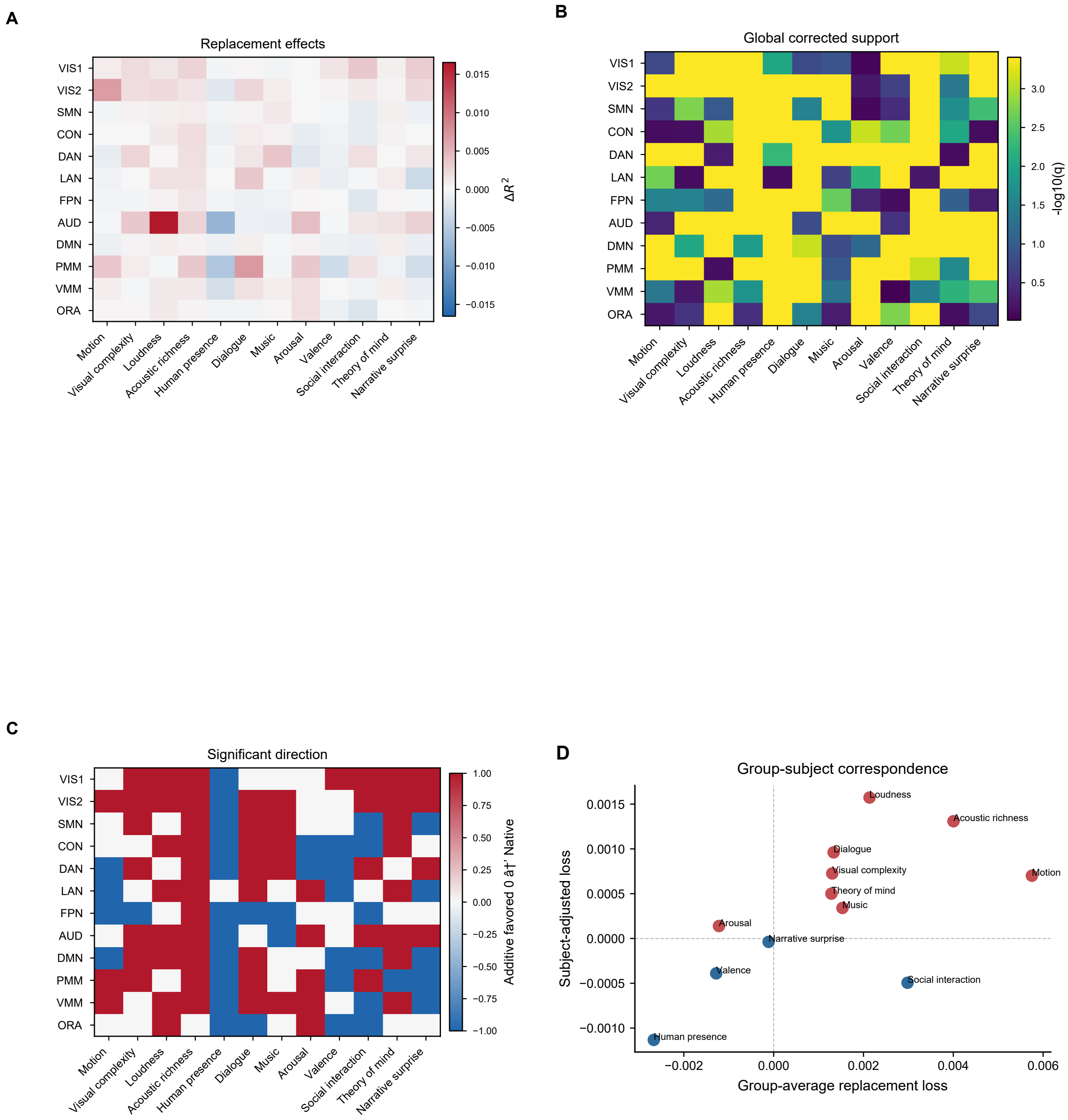


**Supplementary Figure 9 | Full feature-replacement inference.** (A) feature-by-network replacement loss in ΔR²; positive red values indicate information uniquely retained by Native scores and negative blue values favor replacement by Matched scores. (B) −log10 q after global correction of all 144 feature-network tests; q=0.05 corresponds to 1.30. (C) direction of globally significant routes, with red denoting Native-specific, blue Additive-favored, and zero non-significant tests. (D) group-average versus subject-adjusted cortical replacement effects; zero lines define direction quadrants and feature labels identify points.

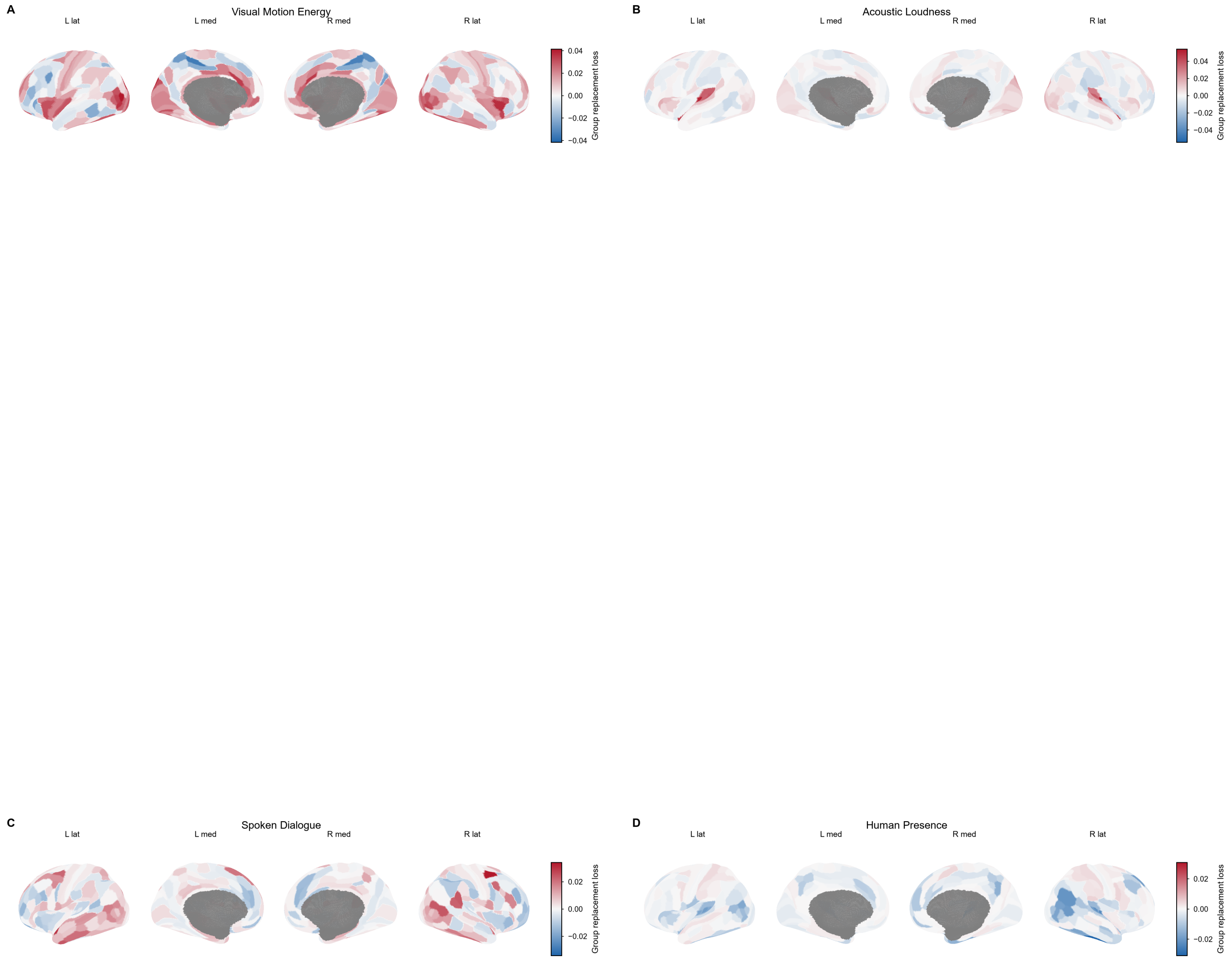


**Supplementary Figure 10 | Representative parcel-level feature-replacement maps.** (A) Visual Motion Energy. (B) Acoustic Loudness. (C) Spoken Dialogue. (D) Human Presence. Values are group replacement loss in $R^2$: positive red cortex indicates prediction loss when the Native feature is replaced by its Matched counterpart, whereas negative blue cortex indicates superior prediction after replacement. Each panel shows four cortical views and uses its own symmetric color limits, reported by the adjacent color bar.

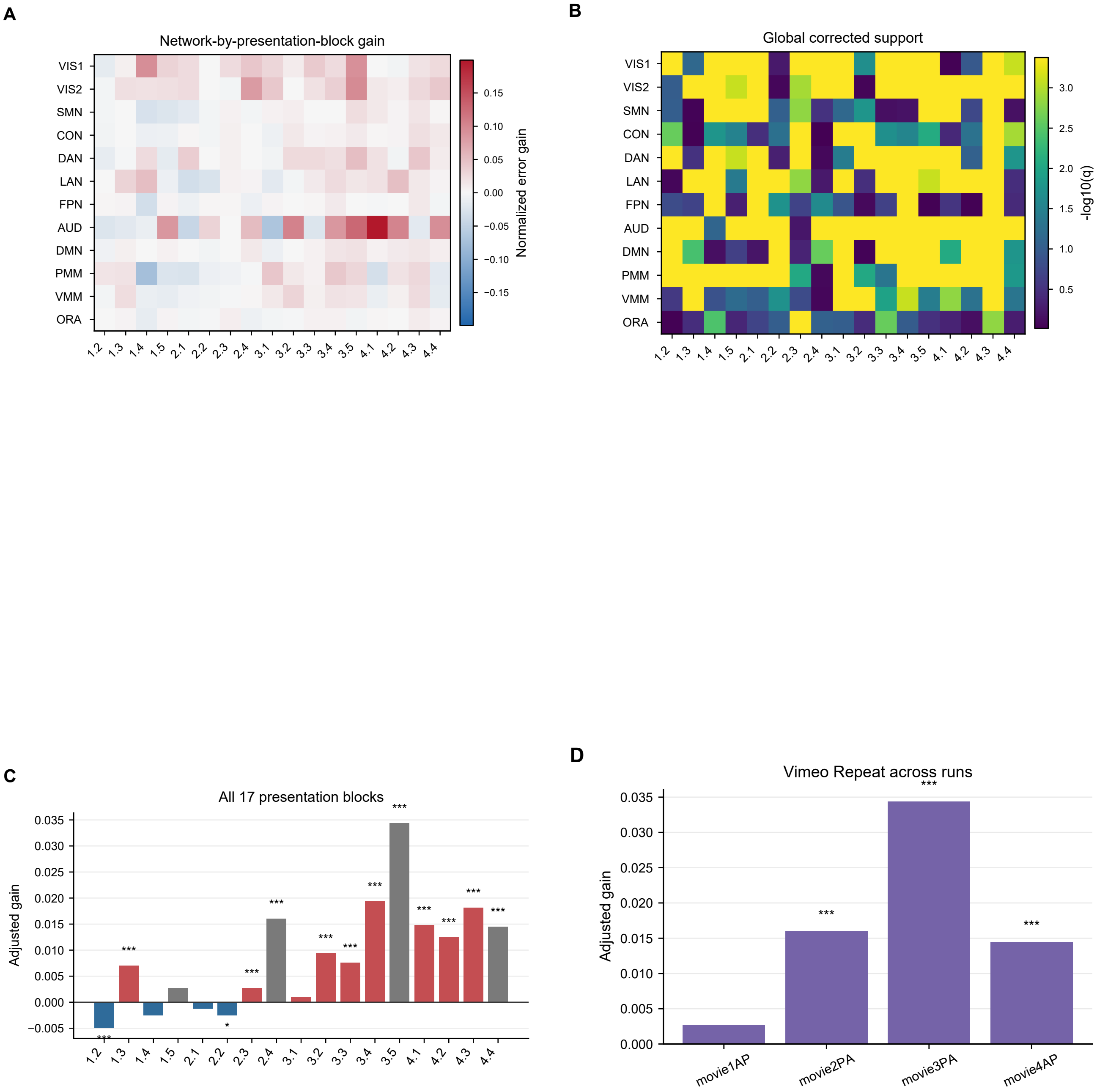


**Supplementary Figure 11 | Source-block and network heterogeneity.** (A) CAB-network by presentation-block matrix of normalized Native error gain; red and blue denote Native- and Additive-favored effects. (B) −log10 q after global correction of all 204 network-block tests; larger values indicate stronger support. (C) adjusted gain for all 17 presentation blocks; grey identifies common-repeat blocks, red and blue show direction for other blocks, the horizontal line denotes zero, and asterisks denote block-wise FDR. (D) adjusted gain for the four Vimeo Repeat presentations; asterisks denote two-sided FDR and bar height is the effect size.

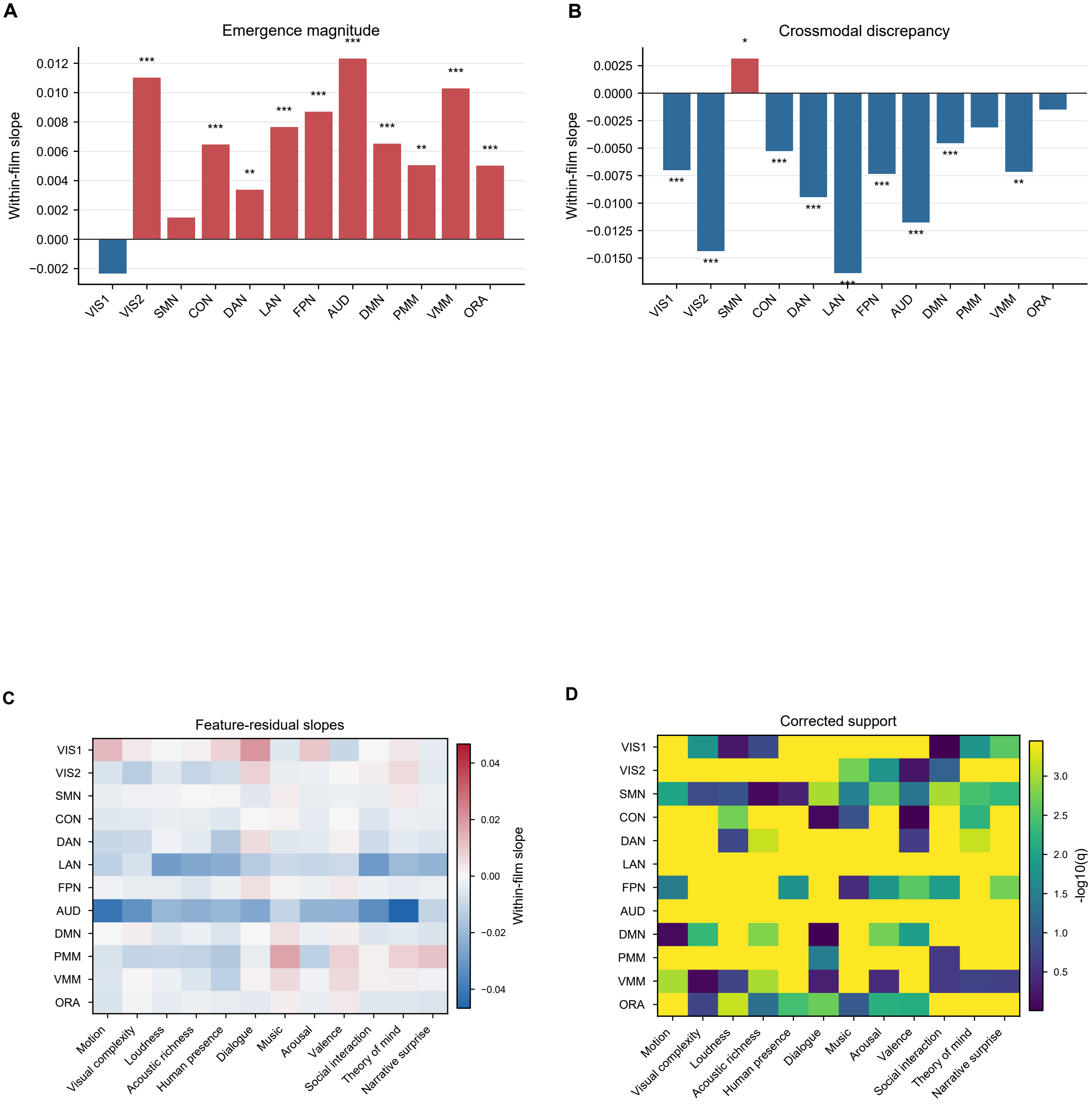


**Supplementary Figure 12 | Complete within-film semantic-association results.** (A) within-film slope relating coherent Native emergence magnitude to network gain. (B) corresponding slope for raw audio-video discrepancy. In A and B, red and blue encode slope direction, the zero line denotes no association, and asterisks denote correction across the 24 prespecified network-mechanism tests. (C) slopes for all 144 feature-residual by network combinations; the diverging color bar gives signed slope magnitude. (D) −log10 globally corrected q for the same tests; q=0.05 corresponds to 1.30.

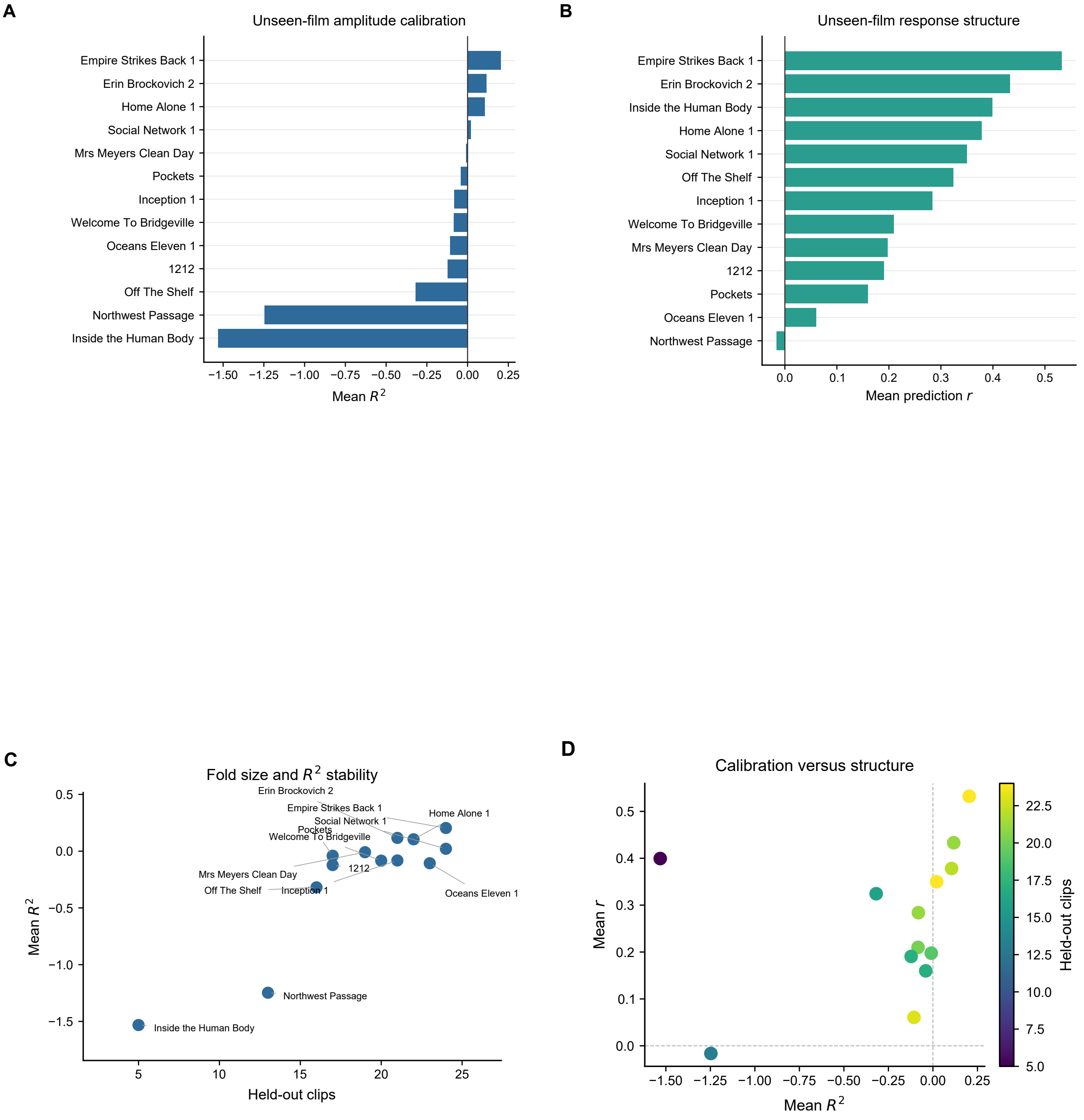


**Supplementary Figure 13 | Leave-one-source-film-out metric audit.** (A) mean fold-wise R² for each completely unseen source film; the zero line separates positive from negative amplitude calibration. (B) mean prediction correlation r for the same folds, quantifying preservation of response structure. (C) held-out clip count versus mean R²; labels identify source films and each point is one fold. (D) mean R² versus mean r; zero lines define calibration/structure quadrants and point color denotes the number of held-out clips.

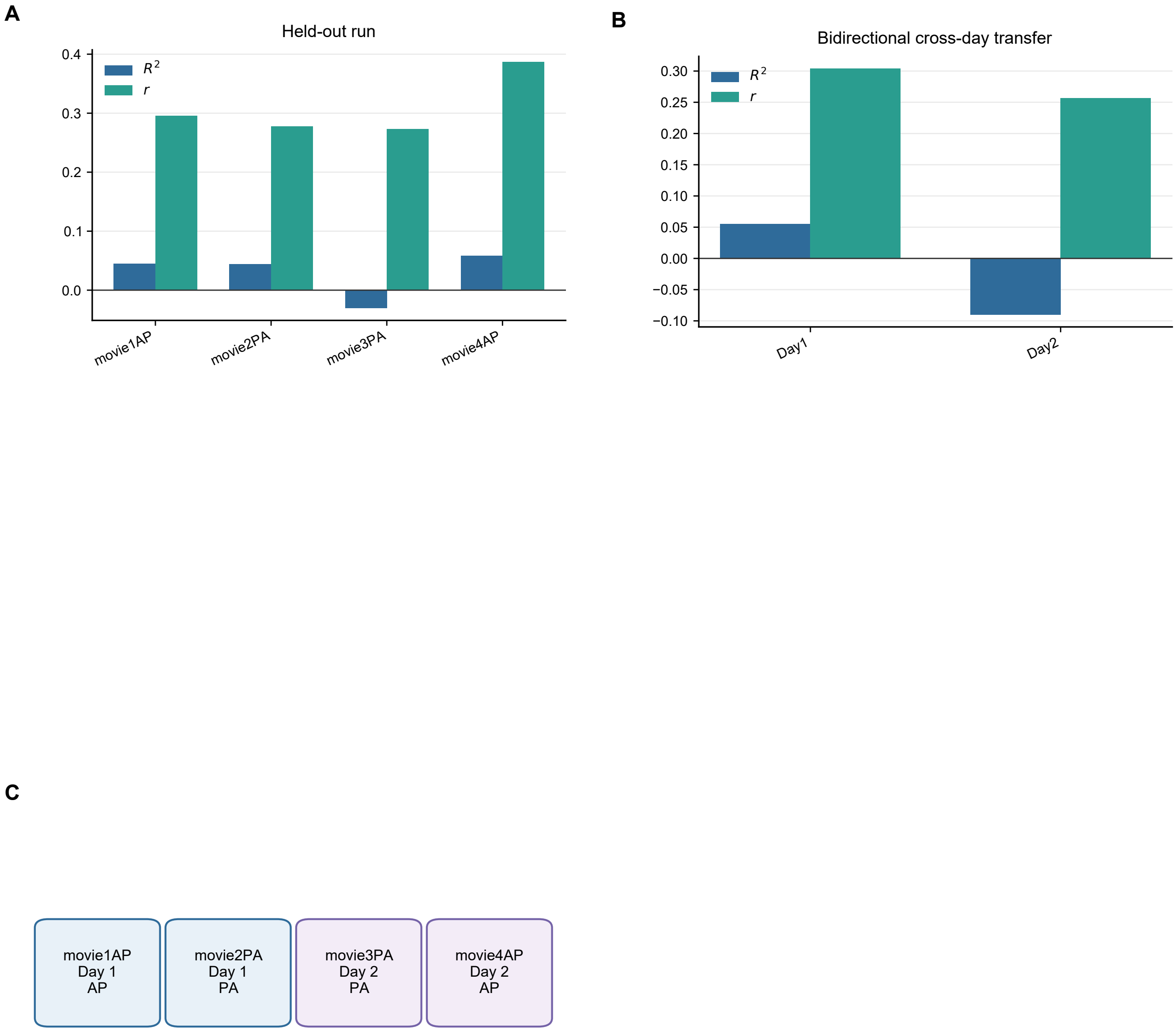


**Supplementary Figure 14 | Unseen-run and cross-day transfer.** (A) mean $R^2$ and prediction r for each held-out movie run; blue and teal bars show the two complementary metrics and the horizontal line denotes zero. (B) bidirectional cross-day transfer using the same metrics. (C) acquisition map linking each movie run to scanning day and AP/PA phase encoding; blue boxes denote day 1 and purple boxes day 2.

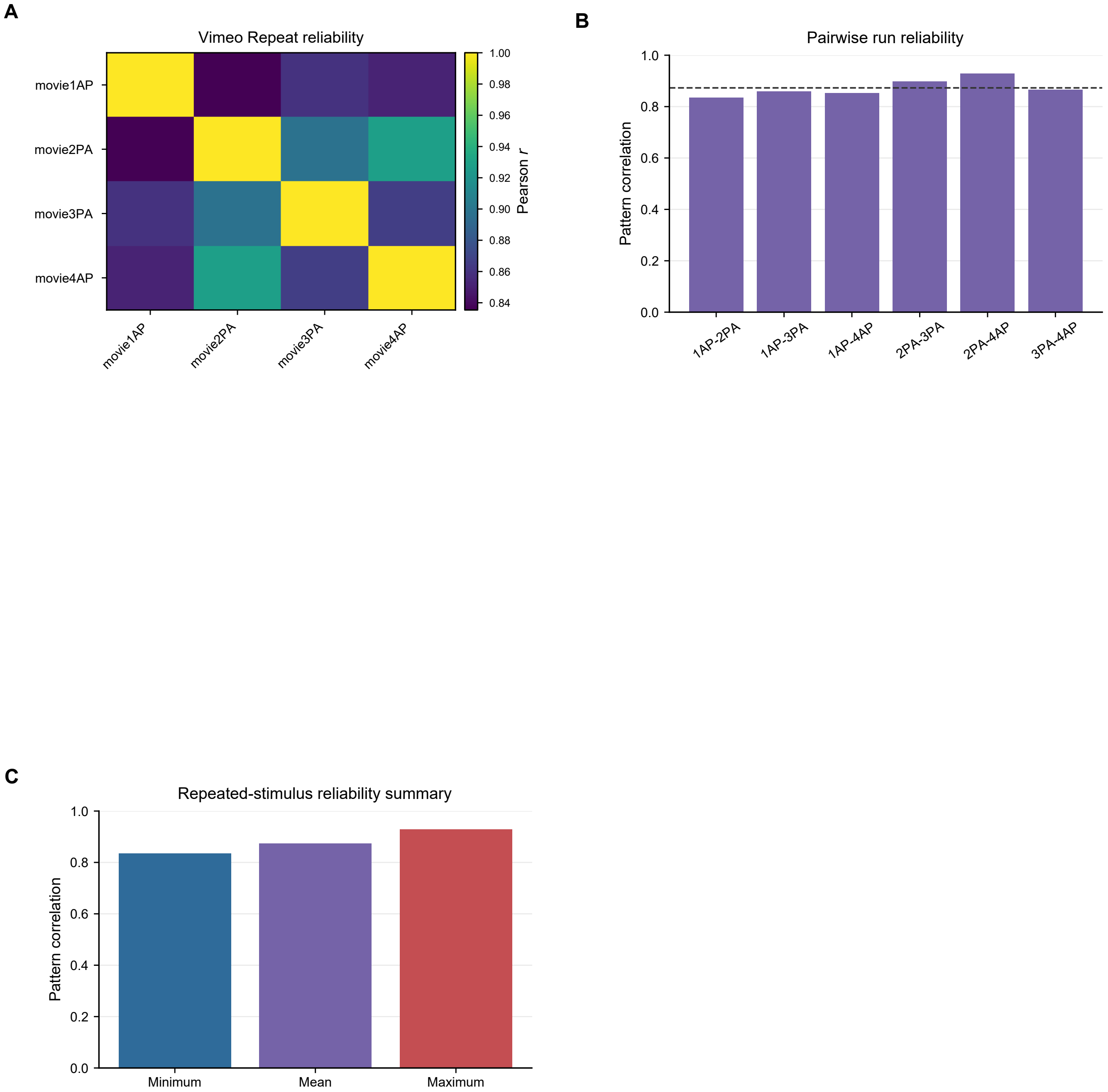


**Supplementary Figure 15 | Repeated-stimulus reliability.** (A) pairwise Pearson correlation matrix of flattened group brain patterns across the four Vimeo Repeat presentations; the sequential color bar spans the observed correlation scale and the diagonal is one by definition. (B) the six unique run-pair correlations; the dashed horizontal line is their mean. (C) minimum, mean, and maximum pairwise correlations, shown on a common 0-1 scale.

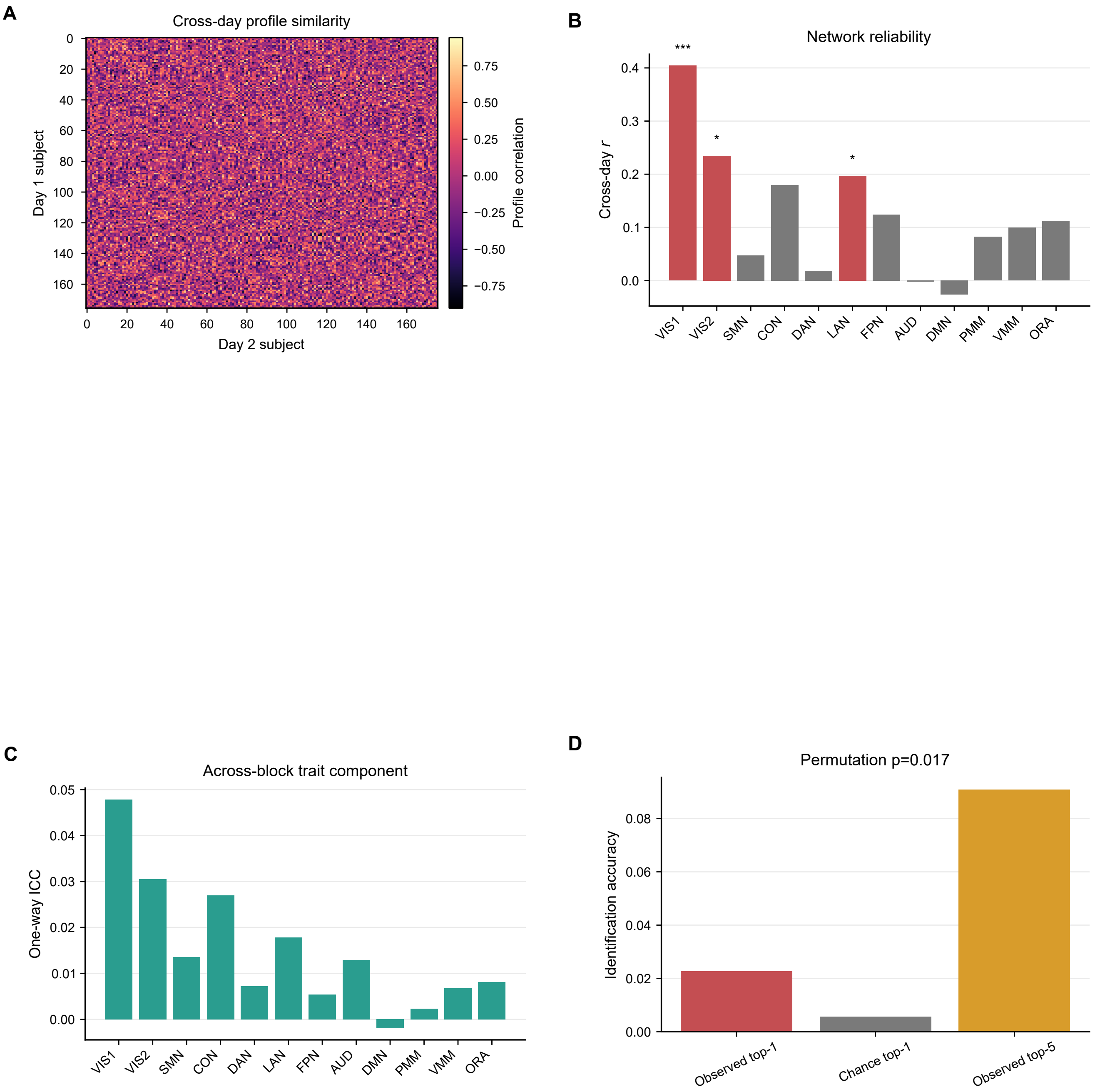


**Supplementary Figure 16 | State-versus-trait analysis.** (A) cross-day subject-profile similarity matrix; rows and columns are day 1 and day 2 subjects and the color bar gives profile correlation. (B) network-wise cross-day r; red bars survive FDR, grey bars do not, and asterisks report corrected q. (C) one-way ICC for each network, quantifying the across-block trait component; bar height is the effect size. (D) observed top-1, chance top-1, and observed top-5 identification accuracy; the title reports the permutation p value.

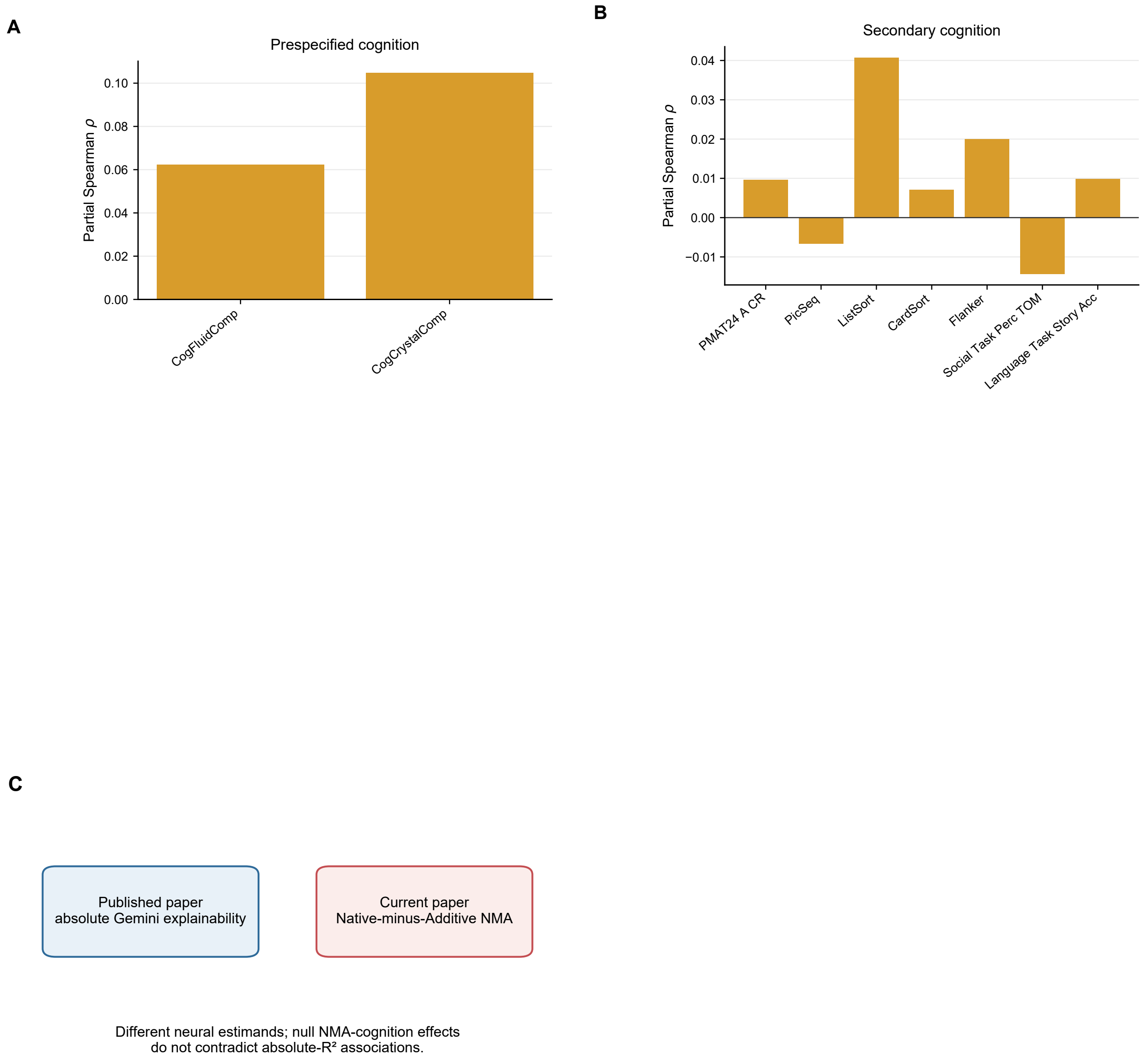


**Supplementary Figure 17 | Cognition.** (A) partial Spearman rho for the two prespecified cognitive outcomes after adjustment for motion, age, and sex; bar height is the association effect size and asterisks denote family-corrected q. (B) corresponding secondary cognitive associations with correction across the secondary family. (C) distinction between the previously published absolute Gemini explainability estimand and the present incremental Native-minus-Additive estimand; the null incremental cognition results therefore do not contradict prior absolute-$R^2$ associations.

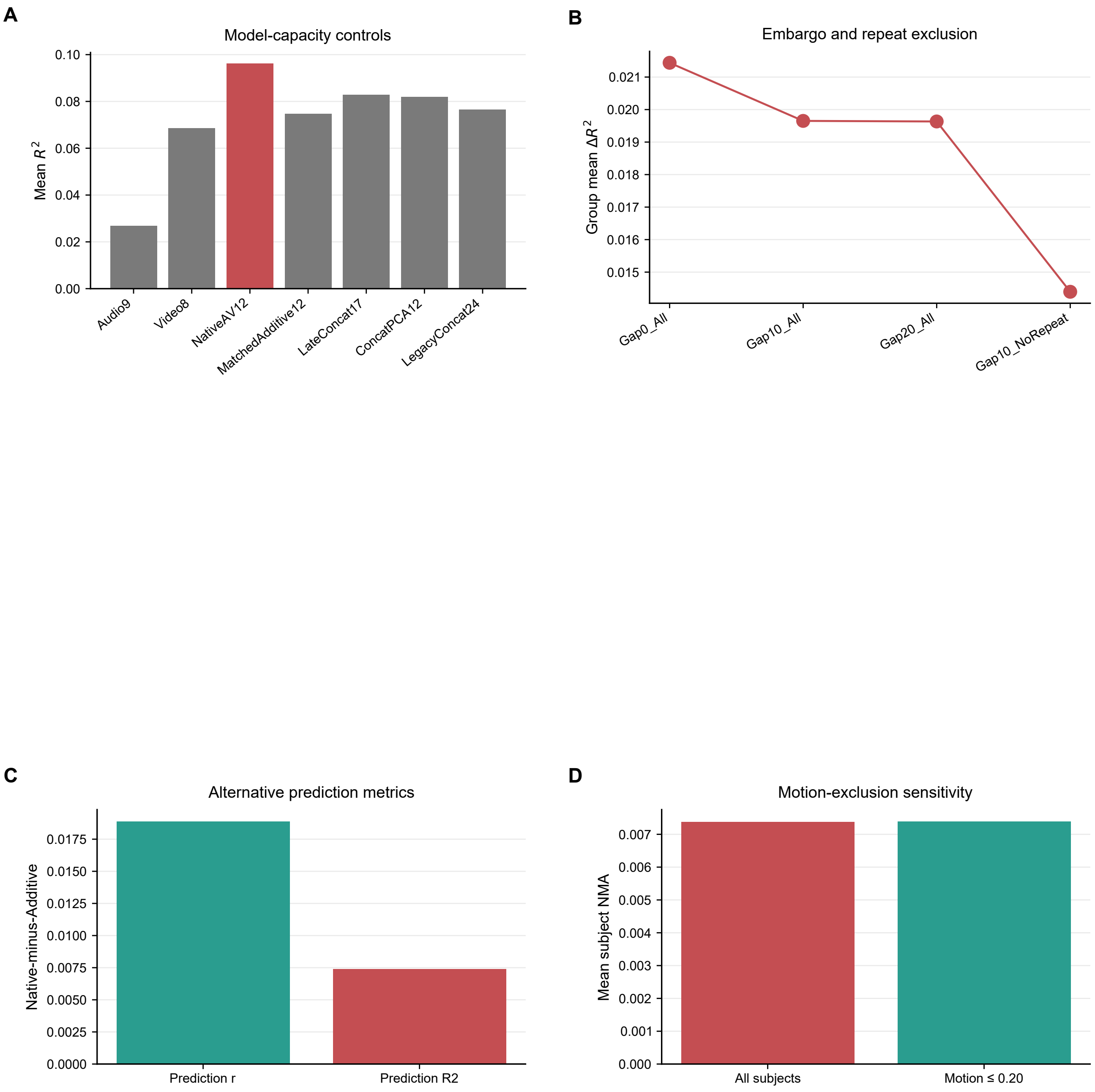


**Supplementary Figure 18 | Additional sensitivity analyses.** (A) mean cross-validated $R^2$ for all model-capacity controls; Native AV is red and alternative models grey. (B) group mean $\Delta R^2$ across temporal embargo and repeat-exclusion conditions; connected points show effect retention. (C) Native-minus-Additive effects expressed as prediction correlation and $R^2$; the zero line denotes equality and bar height is the metric-specific effect size. (D) mean subject NMA in all subjects and after excluding mean framewise motion greater than 0.20; bar height is the sensitivity effect and the plotted source data report sample size.

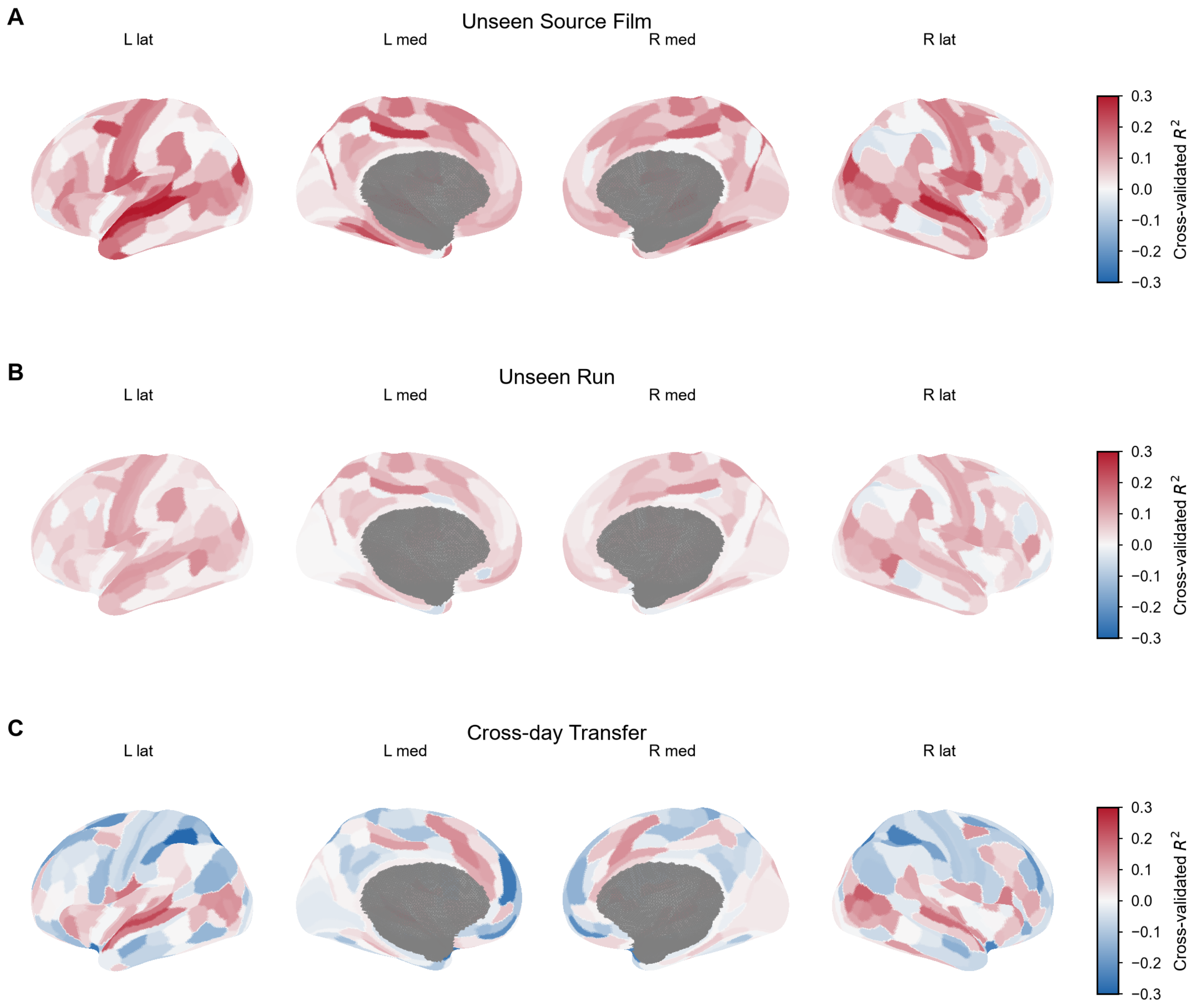


**Supplementary Figure 19 | Cortical topography of absolute amplitude calibration across generalization boundaries.** (A) Unseen source film cross-validated $R^2$. (B) Unseen run cross-validated $R^2$. (C) Cross-day transfer cross-validated $R^2$. All panels show left lateral, left medial, right medial, and right lateral views on the identical −0.30 to 0.30 color scale. Blue denotes negative out-of-sample $R^2$ (amplitude calibration failure), white denotes zero, and red denotes positive $R^2$.
